# Modular safe-harbor transgene insertion (MosTI) for targeted single-copy and extrachromosomal array integration in *C. elegans*

**DOI:** 10.1101/2022.04.19.488726

**Authors:** Sonia El Mouridi, Faisal Alkhaldi, Christian Frøkjær-Jensen

**Affiliations:** King Abdullah University of Science and Technology (KAUST), Biological and Environmental Sciences and Engineering Division (BESE), Thuwal 23955–6900, Kingdom of Saudi Arabia

## Abstract

Efficient and reproducible transgenesis facilitates and accelerates research using genetic model organisms. Here we describe a <u>mo</u>dular <u>s</u>afe harbor transgene insertion (MosTI) for use in *C. elegans* which improves targeted insertion of single-copy transgenes by homology directed repair and targeted integration of extrachromosomal arrays by non-homologous end-joining. MosTI allows easy conversion between selection markers at insertion site and a collection of universal targeting vectors with commonly used promoters and fluorophores. Insertions are targeted at three permissive safe-harbor intergenic locations and transgenes are reproducibly expressed in somatic and germ cells. Chromosomal integration is mediated by CRISPR/Cas9, and positive selection is based on a set of split markers (*unc-119, hygroR*, and *gfp*) where only animals with chromosomal insertions are rescued, resistant to antibiotics, or fluorescent, respectively. Single-copy insertion is efficient using either constitutive or heat-shock inducible Cas9 expression (25 - 75%) and insertions can be generated from a multiplexed injection mix. Extrachromosomal array integration is also efficient (7 - 44%) at MosTI landing sites or at the endogenous *unc-119* locus. We use short-read sequencing to estimate the plasmid copy numbers for eight integrated arrays (6 to 37 copies) and long-read Nanopore sequencing to determine the structure and size (5.4 Mb) of one array. Using universal targeting vectors, standardized insertion strains, and optimized protocols, it is possible to construct complex transgenic strains which should facilitate the study of increasingly complex biological problems in *C. elegans*.

---

The ability to modify the genome of cells and model organisms with high precision has made great advances in the past decade, primarily due to the ease of genetic engineering with CRISPR/Cas9 (Jinek *et al*. 2012; Cong *et al*. 2013; Mali *et al*. 2013). In *C. elegans*, early experiments demonstrated that Cas9 can be expressed from plasmids (Friedland *et al*. 2013; Tzur *et al*. 2013; Waaijers *et al*. 2013; Chen *et al*. 2013; Dickinson *et al*. 2013), mRNA (Lo *et al*. 2013; Chiu *et al*. 2013; Katic and Großhans 2013), or injected directly as a protein (Cho *et al*. 2013; Paix *et al*. 2015) and that templated repair is efficiently mediated from injected plasmids. Subsequent improvements have led to the use of short repair homology (Paix *et al*. 2014), partially single-stranded repair templates (Dokshin *et al*. 2018), improved sgRNA design (Farboud and Meyer 2015), and novel genetic (e.g., Kim *et al*. 2014; Arribere *et al*. 2014; Ward 2015; Farboud *et al*. 2019) and transgenic selection strategies (e.g., Dickinson *et al*. 2015; Schwartz and Jorgensen 2016). These advances have led to the routine use of CRISPR/Cas gene editing to create custom alleles and tagging genes at their endogenous loci.

However, in some cases the goal is to create stable expression of transgenes to determine the effect of regulatory elements on gene expression (Merritt *et al*. 2008), visualize cell populations (Yemini *et al*. 2021), measure cell activity (Kerr *et al*. 2000), or mark genomic regions (Frøkjær-Jensen *et al*. 2014). Initial transgenesis in *C. elegans* was based on transgene expression from extrachromosomal arrays (Stinchcomb *et al*. 1985; Mello *et al*. 1991; Mello and Fire 1995). Extra-chromosomal arrays are formed by injecting linear or supercoiled DNA into the germline syncytium resulting in large hereditary DNA structures that contain many copies of the injected DNA (Stinchcomb *et al*. 1985) propagated as one or more ring chromosomes (Woglar *et al*. 2020). Arrays have the advantage that they are easy to generate and can drive high transgene expression in somatic cells but have the disadvantages that inheritance is unstable with loss occurring during mitosis (Lin *et al*. 2021), expression is mosaic (Okkema *et al*. 1993), and arrays are frequently silenced in germ cells (Kelly *et al*. 1997). Some of these problems can be solved by integrating arrays into the genome by X-ray or gamma irradiation but integration is inherently mutagenic, labor intensive, and does not fully solve the problem of variable expression (Mello and Fire 1995). More recently, alternative methods for integrating arrays have been developed: reactive oxygen species generated by high-intensity blue light can trigger random array integration (Noma and Jin 2018) and Cas9 can be used for targeted array integration into the *dpy-3* or *ben-1* loci (Yoshina *et al*. 2016). However, neither methodology has become widely adopted, possible due to the requirement for a specialized light-source or the mutant phenotype of integrated strains, respectively.

In some cases, expression at near-endogenous levels or expression in the germline is desired, which can be achieved by reducing the number of plasmids. Low-copy number plasmid integrations are primarily generated by biolistic transformation or by using a Mos1 transposon. Biolistic transformation works by bombarding animals with DNA-coated particles, a method that is scalable and can overcome germline silencing (Praitis *et al*. 2001). Biolistic transformation has the disadvantages that a variable number of plasmids are integrated, the insertion site is difficult to determine, and integration may results in chromosomal aberrations (Tyson *et al*. 2018). Targeted single-copy insertions can be generated based on the exogenous Mos1 DNA transposon from *Drosophila* which can transpose in *C. elegans* (Bessereau *et al*. 2001; Robert and Bessereau 2007; Vallin *et al*. 2012). The first widely used single-copy insertion method, Mos1-mediated single-copy insertion (MosSCI), relied on excising transposons in neutral genomic environments and transgene insertion by homology-directed repair from an extra-chromosomal array (Frøkjær-Jensen *et al*. 2008; Frøkjær-Jensen 2015). Biolistic insertion and MosSCI were developed using *unc-119* selection (Maduro and Pilgrim 1995) and subsequently expanded to antibiotic resistance markers (Giordano-Santini *et al*. 2010; Semple *et al*. 2010; Radman *et al*. 2013). Later iterations of Mos1-mediated transgenesis included additional insertion sites and negative selection (Frøkjær-Jensen *et al*. 2012), and random single-copy insertion by transposition enabling a set of universal MosSCI landing sites (Frøkjær-Jensen *et al*. 2014). Single-copy transgene insertions only rely on double-strand DNA breaks and can, therefore, also be generated by CRISPR/Cas9 (Dickinson *et al*. 2013). More recent iterations of single-copy insertion techniques have incorporated specialized landing sites that contain promoters (Silva-García *et al*. 2019), self-selection based on a split antibiotic selection marker (Stevenson *et al*. 2020), or recombinase-mediated cassette exchange (Nonet 2020).

Each of the above methods have their advantages and several are continuously improved (El Mouridi *et al*. 2020; Nonet 2021) yet there is still considerable room for improvement. For example, CRISPR/Cas9-based insertion of transgenes requires different targeting vectors for every location and rely on selection protocols with substantial screening (Dickinson *et al*. 2013). The universal MosSCI sites have the disadvantage that landing sites were generated at random, and the insertion frequency is lower (Frøkjær-Jensen *et al*. 2014). Positive and negative selection markers facilitate the identification of insertions but still require a considerable amount of manual screening and rarely completely avoid false positives (McDiarmid *et al*. 2020; El Mouridi *et al*. 2021). The split antibiotic selection marker facilitates screening but is only compatible with a single antibiotic and genomic location (Stevenson *et al*. 2020). Frequently, it is also not possible to excise the selection marker used to insert transgenes which limits the complexity of transgenic strains that can be generated.

Here, we develop a method we have named <u>Mo</u>dular <u>s</u>afe-harbor <u>T</u>ransgene Insertion (“MosTI”) that improves single-copy and array integration by synthesizing many incremental improvements developed across laboratories and by developing a collection of standardized and validated reagents. MosTI is based on identifying safe-harbor insertion sites located at autosome centers in intergenic regions characterized by permissive chromatin modifications (Ho *et al*. 2014). The method was designed to be highly modular: insertion sites can be rapidly generated and existing sites are easily converted to a different selection marker. MosTI landing sites are compatible with a collection of universal targeting vectors that contain common promoters and fluorophores. The insertion sites were engineered to use a set of split selection markers that obviate the need for negative selection markers as only insertions result in rescue (*unc-119*), fluorescence (*gfp*), or antibiotic resistance (*HygroR*). Finally, MosTI is compatible with both single-copy transgene and targeted array insertion at high frequency. We have validated and characterized several landing sites and developed protocols that allow many single-copy insertions by heat-shocking animals from a single injection. Finally, we use MosTI to resolve plasmid copy-number in integrated arrays and the detailed structure of one array by long-read sequencing.

## Results

### Engineered landing sites and split selection markers enable easy identification of insertions

We set out to develop a modular transgenesis system where insertions are easily identified, new insertion sites can be readily generated, and the selection scheme can be modified. Furthermore, we wanted the ability to integrate single-copy transgenes and extra-chromosomal arrays at well-defined locations. As a first step, we developed a strategy for inserting single-copy transgenes into “safe-harbor” landing sites based on rescuing *unc119*(*ed3*) animals using a split *cbr-unc-119*(+) selection scheme (**Figure 1A**), an approach that is similar to the split antibiotic selection used by (Stevenson *et al*. 2020). The first part of the rescue marker is at the landing site and the second part is located on the target plasmid adjacent to the transgene; neither part can in isolation rescue the *unc-119* phenotype. Injecting plasmids expressing Cas9 and an efficient sgRNA (Moreno-Mateos *et al*. 2015) generates a DNA double-strand break (DSB) at the landing site. The target plasmid contains homology to the DSB site, and homology-directed repair reconstitutes a full rescuing *cbr-unc-119(+)* marker, while also inserting a single copy of the transgene. If desired, the reconstituted split *cbr*-*unc-119* selection marker can be excised after transgene insertion by expressing Cre recombinase. This approach greatly facilitates identification of single-copy insertions: we have observed no false positives and just a few rescued animals are easily visible among a population of Unc animals (see next section). Although not required, we typically enrich for non-rescued transgenic animals with extrachromosomal arrays using fluorescent markers and hygromycin selection.

**Figure 1.**
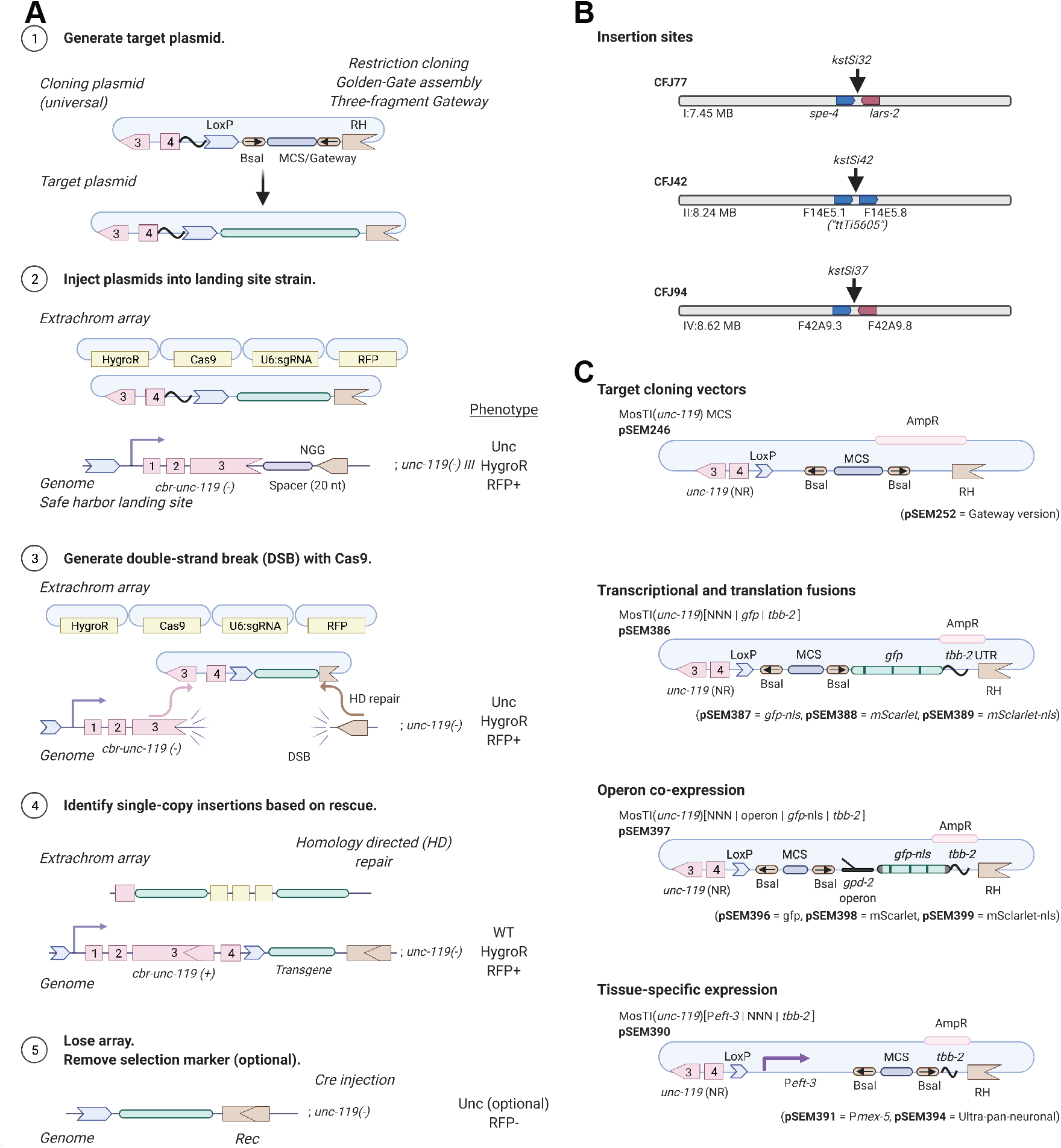
| Schematic of Modular Safe-harbor Transgene Insertion (MosTI) protocol **A**. 1. Target transgenes are generated by cloning (standard restriction-enzyme, Gateway, Golden Gate, or gene synthesis) into a MosTI-compatible plas*F*m*ig*id*ur*c*e*on*1*taining a non-rescuing *cbr-unc-119* rescue fragment. 2. Target plasmids are co-injected with Cas9 and sgRNA plasmids into *unc-19(ed3)* animals containing a MosTI landing site. Antibiotic or fluorescent markers can be used to enrich the population for non-rescued transgenic array animals (optional). 3. Cas9-induced double-strand breaks are repaired by homology-directed repair from the target plasmid, leading to transgene insertion and reconstitution of a functional *cbr-unc-119(+)* gene. 4. Animals with single-copy insertions are identified by phenotypic rescue (N2-like animals on plates with Unc animals). 5. The extrachromosomal array is lost, and the *cbr-unc-119* selection can be removed by Cre expression (optional). **B**. MosTI-compatible insertion sites in safe-harbor landing sites were selected for permissive chromatin environment (Ho *et al*. 2014). **C**. A collection of MosTI-compatible cloning vectors for tissue-specific gene expression, transcriptional and translational fluorescence expression, and fluorescence co-expression with a *gpd-2* operon has been deposited with Addgene (see **Supplementary Figure 3** for all plasmids).

The use of engineered landing sites is useful for several reasons. First, synthetic landing sites allowed us to generate three MosTI landing sites on different autosomes that are compatible with the same target vector and sgRNA (**Figure 1B**). One landing site is near the commonly used *ttTi5605* site on Chr. II (Frøkjær-Jensen et al., 2008), whereas the other sites were chosen for permissive chromatin marks (Ho *et al*. 2014) and minimal disruption to endogenous genes. We inserted landing site in a two-step process: an initial landing site was inserted using a short synthetic *unc-119(+)* selectable marker flanked by LoxP sites; a second injection removed the selection marker by expressing Cre recombinase, leaving behind the final landing site with the split *cbr-unc-119* selection (**Supplementary Figure S1**). This two-step process is efficient and transgenes to generate novel landing sites are easily generated in a one-pot Golden-Gate reaction with a synthetic gene fragment (Engler *et al*. 2008) (**Supplementary Figure S2**). Second, the engineered landing sites are all compatible with a single targeting plasmid. This allowed us to generate standard target vectors compatible with restriction cloning, Golden-Gate cloning, and three-fragment Gateway cloning (**Figure 1C**). We also generated an expanded set of MosTI-compatible cloning vectors: four target vectors with fluorophores (*gfp* and *mScarlet)* for transcriptional and translational fusions (El Mouridi *et al*. 2017), four vectors with a *gpd-2*::fluorophore cassette for transgene co-expression, and three vectors with promoters for ubiquitous (P*eft-3*) (Wheeler *et al*. 2016), germline (P*mex-5*) (Zeiser *et al*. 2011), and neuronal (ultra-pan-neuronal, UPN promoter) (Yemini *et al*. 2021) expression (**Figure 1C** and **Supplementary Figure S3**). We performed whole-genome sequencing on two injection strains (Chr. I: CFJ77 and Chr. II: CFJ42) to identify possible passenger mutations present in the initial *unc-119*(*ed3*) strain or generated by landing site insertion (**Supplementary Table 1**). All plasmids are available from Addgene, and the universal target vector has been deposited with a gene synthesis company for direct gene synthesis instead of standard cloning, which is becoming increasingly convenient and cost-efficient.

We tested MosTI efficiency for single-copy insertions of a plasmid encoding a ubiquitously expressed GFP (P*eft-3*::*gfp*). We chose the *eft-3* promoter because single-copy insertions are bright in somatic cells whereas germline expression is sensitive to silencing in repressive chromatin domains (Frøkjær-Jensen *et al*. 2016). We generated single-copy insertions using constitutive Cas9 expression (P*smu-2::Cas9*) at relatively high frequencies (25-75%)(**Figure 2A**). We also inserted a different transgene (P*rpl-7*::*gfp*) by inducible expression of Cas9 (P*hsp-16.21::Cas9*) using an improved heat-shock protocol (Nonet 2020). These efficiencies are similar to MosSCI and other single-copy transgenesis methods (Frøkjær-Jensen *et al*. 2012; Dickinson *et al*. 2013; Nonet 2020) but do not capture the ease of identifying transgene insertions: we readily observed *unc-119* rescued animals in the F2 or F3 generation (for both protocols) or for many generations after injection using the heat-shock protocol. All rescued animals behaved as expected: the inserted fluorescent marker was expressed, transgenic animals could be rendered homozygous, the extra-chromosomal array was rapidly lost when animals were grown without selection, PCR and Sanger sequencing validated insertion at the intended MosTI landing site, and the *cbr-unc-119* selection marker could be removed by Cre expression without losing the inserted transgene. All P*eft-3::gfp* transgenes inserted at the three MosTI landing sites were expressed at similarly high levels, and all transgenic strains showed GFP expression in the germline (13 of 13 strains), confirming the permissive chromatin environment (**Figure 2B-C**).

**Figure 2.**
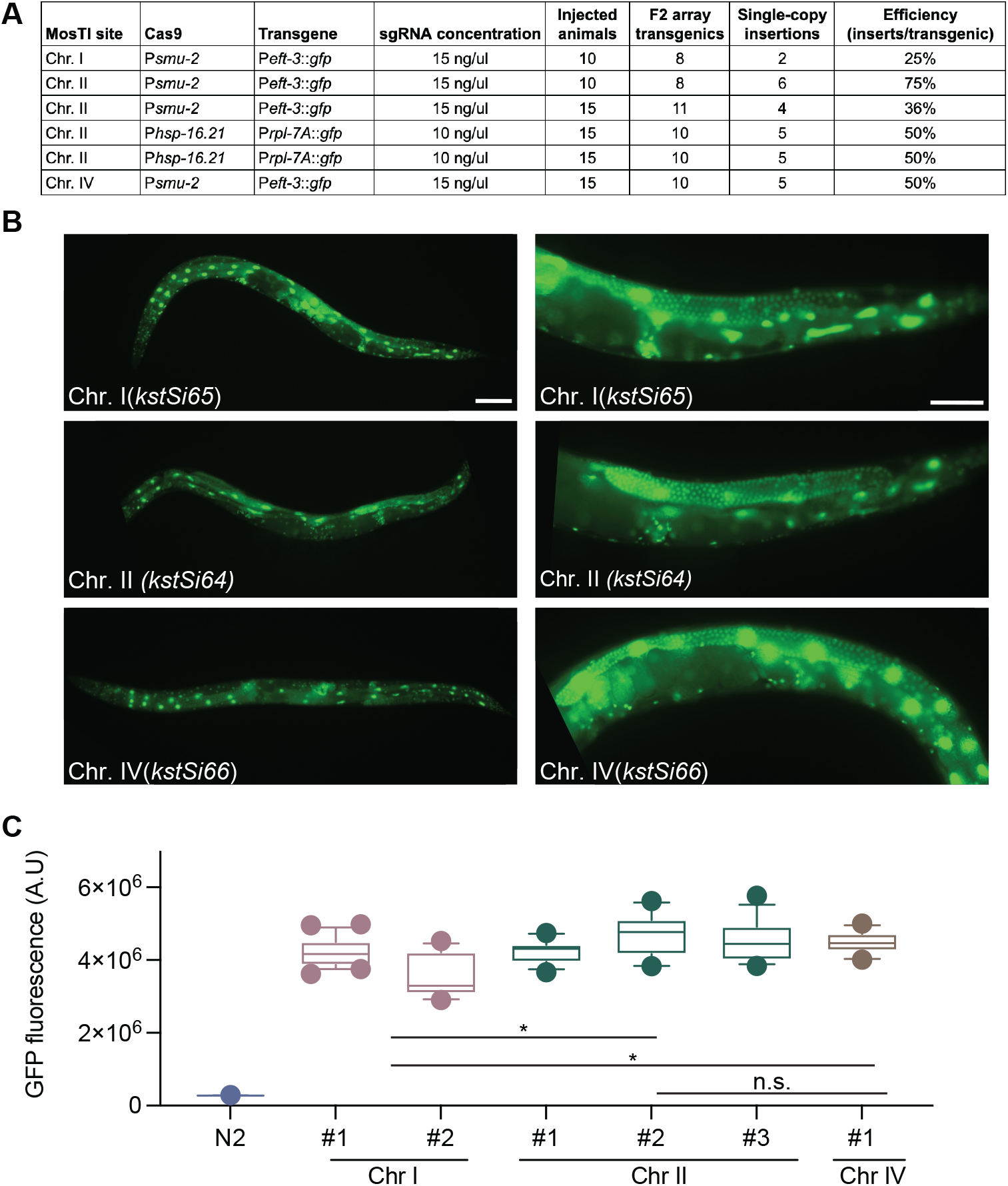
| MosTI insertion frequency and transgene expression **A**. Table with MosTI insertion efficiencies using constitutive (P*smu-2::Cas9*) or heat-shock inducible (P*hsp-16.41::Cas9*) plasmids for Cas9 expression. **B**. Single-copy transgenes (P*eft-3*::*gfp*) are widely expressed, including the germline (13/13 inserts expressed). Left panels 10X magnification, right panels 40X magnification. Scale bars: 20 microns. **C**. GFP quantification by wide-field fluorescence microscopy (L4 stage) from single-copy P*eft-3*::*gfp* insertions at three MosTI landing sites. Statistics: Kruskal-Wallis’s test of fluorescence intensity grouped by insertion site (Chr. I: 35 images, n = 2; Chr. II: 41 images, n = 3; Chr. IV: 13 images, n = 1 insert, * P < 0.05).

The heat-shock protocol (Nonet 2020) requires an additional heat-shock step relative to constitutive expression of Cas9. However, heat-shock induction has the advantage of being highly scalable and requires injection of fewer animals (*i.e*., a few transgenic animals with extrachromosomal arrays are sufficient to generate many independent single-copy insertions). The heat-shock protocol may be particularly useful for the injection of collection of different plasmids simultaneously (“multiplexed MosTI”) and inserting a single copy from the extra-chromosomal array (**Figure 3A**). Conceptually, such an approach could be used to screen many regulatory elements, for example, a library of enhancers or 3’ UTRs coupled to a fluorescent reporter, as previously demonstrated by Kaymak *et al*. (2016). To test the feasibility of multiplexed MosTI, we injected a pool of three target plasmids with visually distinct expression patterns (P*rpl-7A::gfp*, P*eft-3::mMaple*, and P*eft-3::mScarlet*) together with a heat-shock inducible Cas9 plasmid (**Figure 3A**). We established four independent transgenic strains with extra-chromosomal arrays and heat-shocked a single NGM plate of each strain for seven successive generations. In most generations, we were able to generate single-copy insertions form three of the four strains. We generated a total of 29 independent single-copy insertions from these three strains and observed no obvious decline in efficiency over time (**Figure 3B**). In some cases, we were able to isolate animals with every type of transgene insertion from a single NGM plate (**Figure 3C**), demonstrating the feasibility of scaling the approach further. Several observations were notable: first, only one of 29 transgene insertions was non-fluorescent, probably caused by an indel in the transgene, as previously observed (Frøkjær-Jensen *et al*. 2008). Second, one of the four transgenic strains yielded no insertions in any generation. We did not determine the exact cause but speculate that the target spacer at the MosTI site was destroyed by NHEJ in an early generation or that this array was somehow not permissive for Cas9 expression. Third, not all transgenes were inserted at similar frequencies, possibly due to differences between the injected plasmids (the *gfp* plasmid was linearized whereas the *mMaple* and *mScarlet* plasmids were supercoiled). We recently showed that linearized plasmids were incorporated into arrays more efficiently than supercoiled plasmids (Priyadarshini *et al*. 2022); the integration frequency may simply reflect the relative abundance of plasmids in the array. We note that a conceptually similar approach was developed by Kaymak *et al*. (2016) using MosSCI to assay 3’ UTR gene regulation but their approach was not scalable in the same way as MosTI.

**Figure 3.**
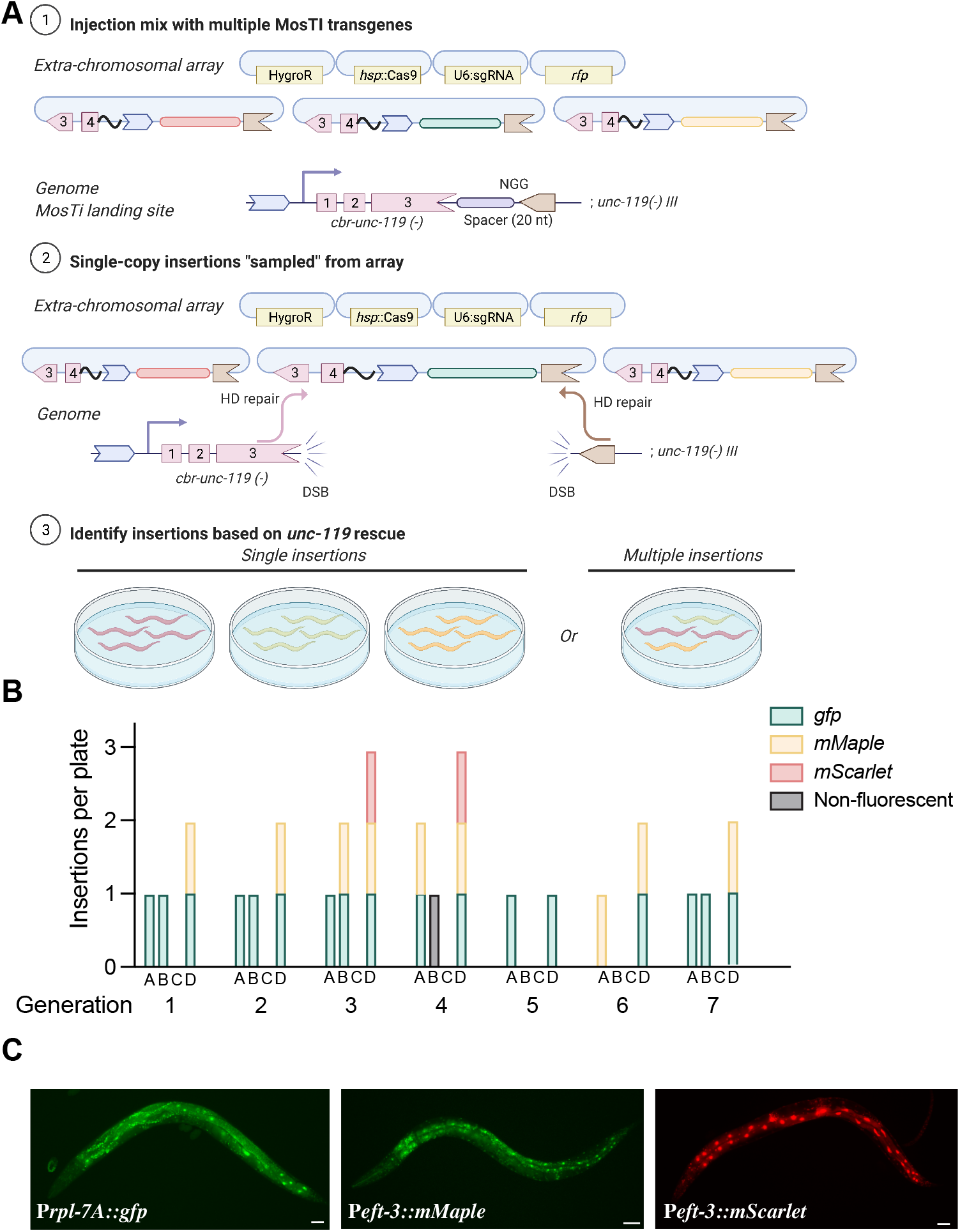
| Single-copy insertions from multiplex transgene injections **A**. Schematic of multiplex insertions using MosTI. Multiple transgenes are pooled in one injection mix. During homology-directed repair, a single transgene from the array is inserted into the MosTI site. **B**. Quantification of single-copy MosTI insertions over seven generations from four independent transgenic animals with extra-chromosomal arrays containing three target transgenes (P*rpl-7A*::*gfp*, P*eft-3*::*mMaple3*, P*eft-3*::*mScarlet*) and a heat-shock inducible Cas9 (P*hsp-16.21*::*Cas9*). In each generation, a new population of animals was generated from a non-rescued (Unc) animal carrying an extra-chromosomal array and these animals were heat-shocked one or two days before exhausting the bacterial lawn. **C**. Fluorescence microscopy showing expression from single-copy insertions of each of the three target transgenes. 20X magnification, scale bar = 20 microns.

In sum, MosTI is an efficient method to insert single-copy transgenes into well-defined genomic locations using several protocols. All landing sites are compatible with robust somatic and germline expression. From a practical perspective, the split selection markers make it easier to isolate animals with single-copy transgenes and the heat-shock protocol is a simple method to generate many inserts from only a few injections.

### Exchanging MosTI selection markers is simple

The use of a strong phenotypic selection marker, such as *unc-119*, has the benefit that insertions are easy to identify, and insertions are easily rendered homozygous. However, *unc-119* animals are slow growing and harder to inject than wild type animals (Maduro and Pilgrim 1995). We designed MosTI to be modular and have developed a standardized strategy to convert selection markers at MosTI landing sites. Conversion is simple: we use MosTI to insert a transgene containing a second split selection marker and a new spacer sequence (**Figure 4A**). In a second (optional) step, the *cbr-unc-119* marker is removed by Cre-mediated recombination. We validated this conversion strategy using a split hygromycin resistance gene, similar to Stevenson *et al*. (Stevenson *et al*. 2020), and a split fluorescent marker that gives bright muscle fluorescence (P*mlc-1*::*gfp*) (El Mouridi *et al*. 2020). We converted MosTI sites to HygroR (Chr. II) and *gfp* (Chr. I and Chr. II) and generated single-copy P*mex-5*::*gfp* insertions (**Figure 4B**). Both selections were efficient and resulted in GFP expression in the germline from the *mex-5* promoter. However, we found that identifying fluorescent animals (four inserts from 15 injected animals) (**Figure 4C**) was considerably easier than isolating animals with antibiotic resistance (one insert from 20 injected animals). MosTI insertions most frequently happen in the F2 generation when food is almost exhausted: it was relatively easy to identify moving (*unc-119* rescued) or fluorescent (*gfp* selection) L1 animals on a starved plate. In contrast, it was harder to identify healthy animals on hygromycin, which is most effective at preventing larval growth and takes several days to kill animals (Radman *et al*. 2013).

**Figure 4.**
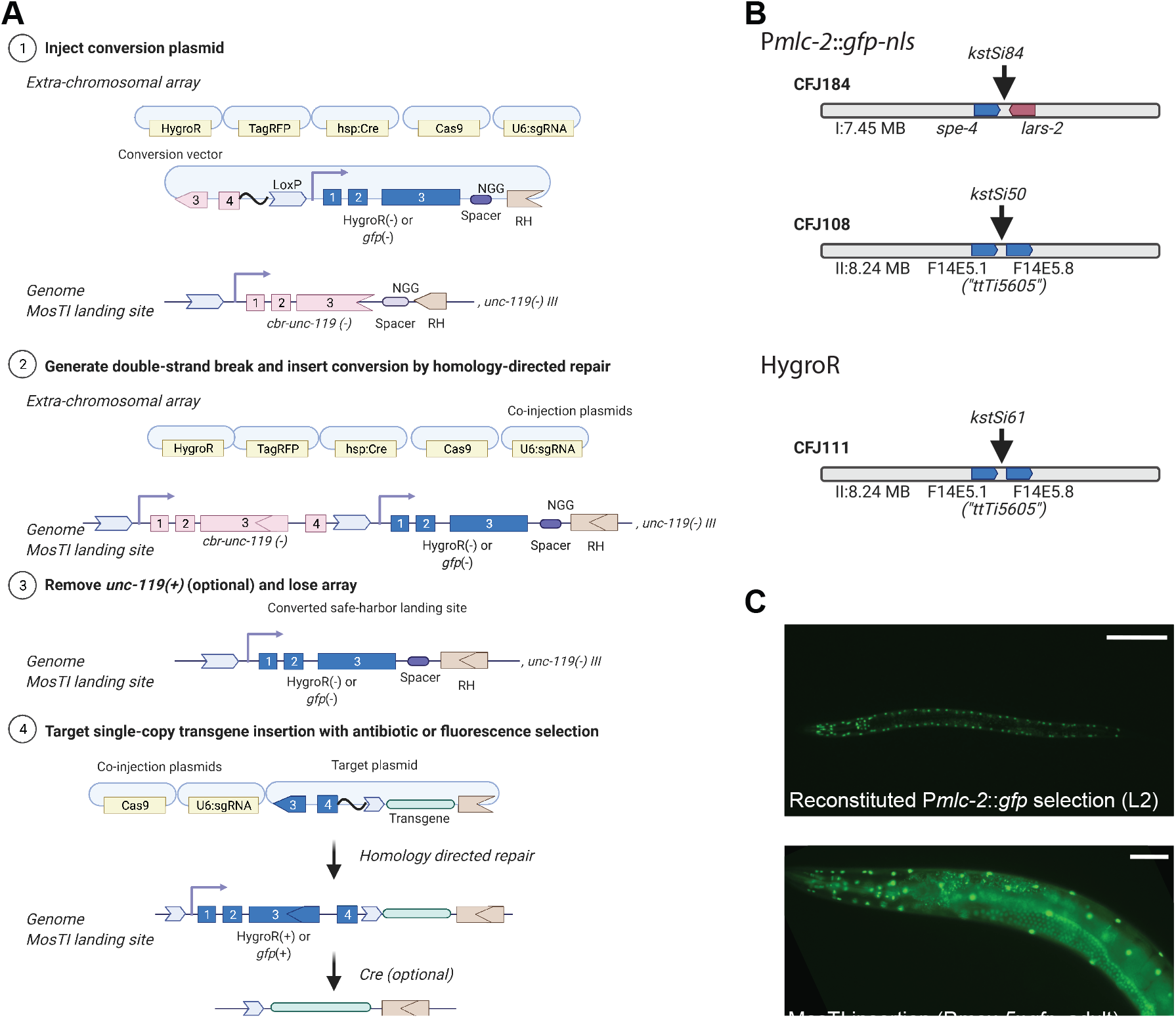
| MosTI sites are modular and can be converted to other split selection markers **A**. Schematic of the conversion of an *unc-119* MosTI site into a MosTI site that uses a different split selection marker (here, HygroR or P*mlc-2*::*gfp*). 1. Injection of a conversion plasmid (*e.g*., generated by gene synthesis) into *unc-119*(*ed3*) mutants with a MosTI landing site. 2. Insertion of the conversion plasmid generates a new MosTI landing site using a different split selection marker. 3. Loss of the extra-chromosomal array and (optional) excision of the reconstituted *cbr-unc-119* selection marker. 4. Injection of a modified MosTI target vector uses the novel selection marker to insert a single-copy transgene. **B**. Available MosTI sites for split P*mlc-2*::*gfp*-*nls* and HygroR selection. **C**. Example of a single-copy insertion of a P*mex-5*::*gfp* (germline) transgene into a converted MosTI site using a split pan-muscular *gfp* selection marker (P*mlc-2*::*gfp*-*nls*). Left panel: Pan-muscular nuclear GFP expression (L2 stage). Right panel: Nuclear GFP expression in muscles (from pan-muscular selection) and in the germline (from the P*mex-5*::*gfp*-*nls*) transgene. Left panel 20X magnification, right panel 40X magnification. Scale bar: 20 microns.

Regardless of the preferred selection, we show that it is relatively easy to change a MosTI site to fluorescence or antibiotic selection. The conversion vector, targeting vectors, and sgRNA can be cheaply generated by gene synthesis with minimal cloning. By modifying existing designs and following a few simple design guidelines, additional safe-harbor landing sites are compatible with the standard collection of targeting vectors. We also note that *cbr*-*unc-119* selection can be “re-used” by inserting a second split *unc-119* fragment. This could, for example, be used to iteratively generate strains with multiple, tightly linked single-copy transgene insertions.

### Split selection markers allow targeted integration of extra-chromosomal arrays

In some cases, simultaneous expression of many transgenes is needed. For example, a method for automated neuron identification relies on expression from 41 different fluorescent promoter constructs (Yemini *et al*. 2021). In other cases, high expression of a single transgene may be necessary, for example, for genetically encoded sensors (e.g., Kerr *et al*. 2000; Flytzanis *et al*. 2014; Hashemi *et al*. 2019) or for spatiotemporal control of gene expression with bi-or tri-partite effectors (Wei *et al*. 2012; Wang *et al*. 2018). Transgene arrays, which contain several hundred plasmids (Stinchcomb *et al*. 1985), are useful for this purpose, especially when arrays are chromosomally integrated to stabilize expression and avoid mitotic loss in cell division (Mello and Fire 1995). We have adapted the split MosTI selection for a CRISPR/Cas9-based method that uses non-homologous end-joining (NHEJ) to target arrays for integration (Yoshina *et al*. 2016; Yoshina and Mitani 2022) (**Figure 5A**). The adaptation required a short non-rescuing *unc-119* fragment (“array target fragment”) to mediate array integration at MosTI landing sites by NHEJ. In this scheme, both the array target fragment and the split *cbr-unc-119* selection marker at the MosTI landing site are cut by the same sgRNA. The DSBs are generated in an intron, so NHEJ between the array and insertion site can reconstitute the full *unc-119* selection marker even if the repair generates short indels (**Figure 5A**). We first tested array integration efficiency at a MosTI site (Chr. II) using a two-step protocol (“two-step plasmid”) which avoided co-integrating the Cas9 and sgRNA plasmids. We first generated transgenic MosTI animals with an extra-chromosomal array containing two fluorescent markers (P*mlc-1::gfp*, P*mlc-1::tagRFP*), a plasmid encoding antibiotic resistance (HygroR), and the array target fragment. Like the protocol for single-copy insertions, these transgenic array animals expressed the injected fluorophores and were resistant to hygromycin but were not rescued for the *unc-119* phenotype. We injected these transgenic animals with a second mix containing plasmids that expressed Cas9 and an sgRNA to generate DSBs. We successfully identified integrated arrays based on *unc-119* rescue at frequencies ranging from 20 to 33% of injected animals (**Figure 5D**). As expected, the rescued animals with the integration expressed all fluorophores from the initial extra-chromosomal array (**Figure 5B**) and could be rendered homozygous. Most strains with array insertions were fully rescued for the Unc-119 phenotype except for one strain with particularly bright expression that showed variable rescue. This integration was likely an example of somatic position effect variegation since both rescued and non-rescued animals segregated a mix of rescued and non-rescued animals over several generations (**Supplementary Figure S4**).

**Figure 5.**
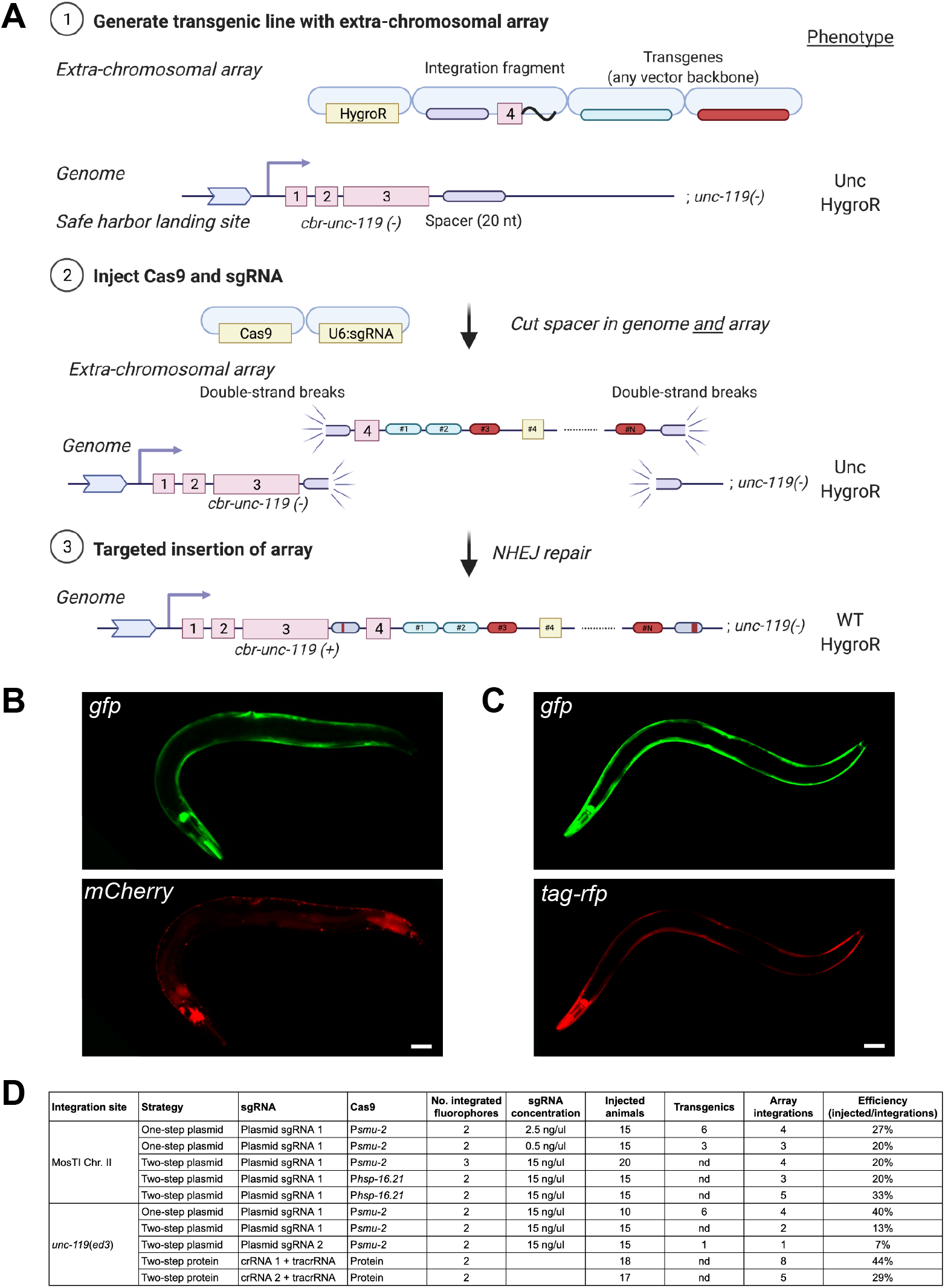
| Targeted integration of extrachromosomal arrays at MosTI sites and at the endogenous *ce-unc-119* locus **A**. Schematic of “two-step plasmid” integration of extrachromosomal arrays at a MosTI landing site. 1. An extrachromosomal array is formed by injecting transgenes, including a plasmid containing the fourth intron of *cbr-unc-119* (“integration fragment”) into *unc-119(ed3)* animals containing a MosTI landing site. 2. Plasmids expressing Cas9 and an sgRNA injected into transgenic animals cause double-strand breaks (DSBs) at the MosTI landing site and in the array (in integration fragments). 3. DSBs are repaired by non-homologous end-joining (NHEJ) between the array and the MosTI site, resulting in *unc-119* rescue. The sgRNA target is in an intron to allow rescue even when NHEJ repair causes short indels. Note, arrays can also be inserted by a similar strategy at the *ce-unc-119(ed3)* locus, or by injecting Cas9 protein and crRNA/tracrRNA. **B**. Fluorescence microscopy of an integrated extra-chromosomal array containing transgenes expressed in muscles (P*mlc-1*::*gfp* and P*rab-3::mCherry*) inserted into a MosTI site on Chr. II. Scale bar = 20 microns. **C**. Fluorescence microscopy of an integrated array (P*mlc-1*::*gfp* and P*mlc-1::tagRFP*) inserted into the endogenous *ce-unc-119(ed3)* locus (Chr. III). Scale bar = 20 microns. **D**. Table showing the efficiency of extra-chromosomal array insertion using one-step and two-step protocols at MosTI landing sites and at *ce-unc-119*(*ed3*).

Although array insertion into a MosTI site may be preferable, the same integration method should work at any genetic locus with a strong loss-of-function phenotype caused by a late mutation. Many laboratories have the commonly used *unc-119(ed3)* strain and we, therefore, tested insertions into the *unc-119* locus using a modified integration fragment and sgRNA. Using the same two-step plasmid protocol, we were able to integrate arrays into the *unc-119* locus at similar frequencies to MosTI sites (7 to 40% efficiency) (**Figure 5C-D**). We tested three additional protocols for generating array insertions. In one protocol (“one-step plasmid”), we injected all plasmids in a single mix (fluorophores, Cas9, sgRNA, and integration fragment). Rescued integrants from this technique were obvious already in the F1 generation and were generally dimmer than integrants from the “two-step plasmid” protocol (see next section). These integrants may correspond to smaller arrays that would not have become stable inherited arrays (Mello *et al*. 1991; Lin *et al*. 2021). In a third protocol, we first generated an intermediary transgenic injection strain with plasmids encoding a heat-shock inducible Cas9 (*hsp-16.2*::*Cas9*) and hygromycin selection. We used this strain for a second injection with array plasmids (fluorophores, integration fragment, and sgRNA) and generated animals with two arrays. We heat-shocked these animals and were able to generate array insertions at reasonably high frequencies (20 to 33%) (**Figure 5D**). Finally, we also tested the initial two-step plasmid protocol using Cas9 protein and pre-complexed crRNA/tracrRNA guides (“two-step protein”) to integrate arrays in *unc-119(ed3)* animals. The protein-based protocol resulted in similar or higher efficiencies depending on the cut-site and crRNA used (29 to 44%)(**Figure 5D**).

In sum, MosTI can be used to target extra-chromosomal arrays for integration using *unc-119* selection and a variety of protocols with similar efficiencies. We have not attempted integrations at MosTI sites using other selection markers, but the approach should work for hygromycin or fluorescence selection. The choice of protocol will depend on the desired size of integrations, how many integrations are needed, and whether continued expression of Cas9 may cause a problem for subsequent experiments. We note that we have not observed arrays changing over time, despite integration of Cas9 and sgRNA plasmid (see next section).

Conceptually similar protocols can generate array integrations at the endogenous *unc-119* locus and can presumably be extended to most other endogenous genes with a strong loss-of-function phenotypes.

### Molecular characterization of targeted array insertions

An early study demonstrated that arrays formed from super-coiled plasmids contain sequences organized in tandem, with some plasmids containing short duplications and deletion (Stinchcomb *et al*. 1985). The size and repetitive nature of simple arrays formed from plasmid DNA preclude full assembly using Illumina short-read sequencing. However, long-read sequencing methods, such as Oxford Nanopore and PacBio sequencing, facilitate assembly of complex and repetitive genomic regions (Logsdon *et al*. 2020). A recent study assembled a complex extra-chromosomal array formed from yeast genomic DNA and linear transgene fragments by Nanopore sequencing (Lin *et al*. 2021). The array was mainly formed by non-homologous end-joining of yeast genomic DNA and spanned 50 contigs with a total estimated length of 11 Mb. In another study, Tyson *et al*. (2018) characterized a low-copy plasmid integration generated by biolistic bombardment using Nanopore sequencing. The biolistic integrant contained 30 kb DNA but none of the transgenes (P*pie-1*::*gfp*) were fully intact and the integration resulted in a 2 Mb duplication of the flanking region.

To better characterize integrated arrays formed from circular plasmid DNA and to detect large-scale genomic changes, we first performed short-read Illumina whole-genome sequencing (WGS). We characterized arrays integrated at the *unc-119* locus by the one-step plasmid protocol (“plasmid-integrated strains”) and the two-step protein method (“protein-integrated strains”). All injections contained supercoiled plasmids encoding two fluorophores (P*mlc-1*::*gfp* and P*mlc-1*::*tagRFP*), the integration fragment, and linear stuffer DNA (1kb plus DNA ladder, Invitrogen). Injections with Cas9 plasmid were super-coiled (P*smu-2::cas9::mCherry*) and protein strains contained a supercoiled plasmid encoding an antibiotic resistance marker (HygroR) (**Figure 6A**).

**Figure 6.**
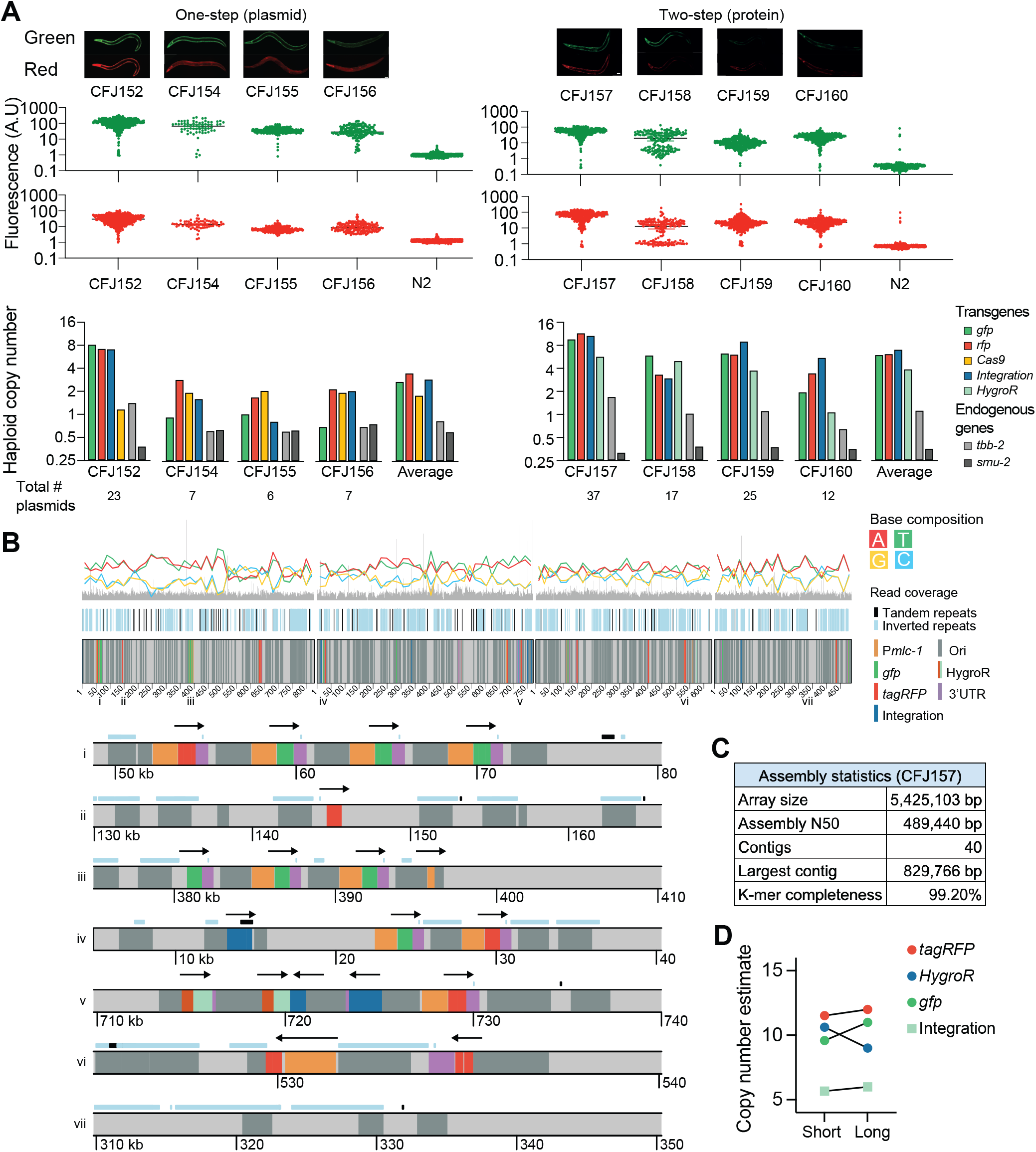
| Molecular characterization of targeted array integrations **A**. Analysis of arrays integrated at the *ce-unc-119* locus using a “one-step plasmid” protocol (left) and a “two-step protein” protocol (right). Top: Fluorescence images of strains with array integrations. Scale bar = 20 microns. Middle: Quantification of total fluorescence in adult animals using a COPAS large-particle fluorescence sorter. Bottom: Plasmid copy-number estimates based on whole-genome sequencing (Illumina short read sequencing). **B**. Integrated array assembly from long-read Oxford Nanopore sequencing of CFJ157 (two-step protein array integration containing *gfp, tagRFP, hygroR*, integration fragment, and 1 kb plus ladder). Top: Schematic overview of four long contigs. Bottom: Individual areas from contigs (location indicated by numbers above). **C**. Assembly statistics from the strain CFJ157. **D**. Comparison of copy-number estimates based on Illumina short-read sequencing and Oxford Nanopore long-read sequencing of CFJ157.

We used WGS data to estimate the copy number of all plasmids in the injection mix based on average read coverage (**Figure 6A**). We detected at least one copy of all injected plasmid (from two to 23 copies) in every strain and the integrated arrays contained between six and 37 plasmids in total (**Figure 6A**). Our plasmid copy-number estimates are likely correct within a factor of two: our analysis estimated the copy number of two endogenous genes to 0.97 ± 0.14 for *tbb-2* (47% GC) and 0.48 ± 0.06 for *smu-2* (30% GC). Plasmid-integrated strains were less bright compared to protein-integrated strains, and this correlated with lower estimated plasmid copy-number (**Figure 6A**). All arrays contained multiple copies of the integration fragment with two strains (one plasmid-integrated and one protein-integrated) containing intact spacer sequences, showing that Cas9 cutting efficiency was sometimes incomplete using either integration protocol (**Supplementary Figure S5**). A couple of observations are worth noting: in the plasmid-based protocol, both sgRNA and a germline-expressed Cas9 are integrated which could possibly delete parts of the array over time. However, we propagated a plasmid-integrated strain with several intact copies of the integration plasmid for ten generations and observed no change in fluorescence (**Supplementary Figure S6**). Also, all four protein strains were derived from a single transgenic extra-chromosomal array strain. Given the variation in plasmid-copy number it is likely that only some fragments of the original array were integrated. In this model, Cas9 protein injection “catastrophically” cuts the array into fragments at most spacer sequences and the genomic location; NHEJ between array fragments and the genomic locus results in partial array integration. If this model is correct, then the relative copy-number of the integration fragment in the injection mix could be used to tune the size of the integrated array.

The *unc-119* selection scheme tolerates indels at the cut-site because the spacer is in an intron. However, large indels or duplications near the cut-site could be problematic by perturbing nearby gene expression. We analyzed the WGS data to detect small and large-scale indels near the insertion site based on read coverage (**Supplementary Figure 7**). None of the integrated strains contained short or large-scale deletions and we found only one instance of a putative short duplication (approximately 6 - 7 kb of *unc-119*) (**Supplementary Figure 7**).

To determine the structure of a simple array (*i.e*., an array composed mainly of plasmid DNA and repetitive DNA ladder) we performed long-read Oxford Nanopore MinION sequencing on a strain with an integrated array containing a relatively high copy number (CFJ157). We were able to assemble the array into 40 contigs with an N50 value of 489 kb. We estimated the total array size at 5.5 Mb (**Figure 6C**), which is within the range of estimates based on microscopy (1 to 2.2 Mb) (Woglar *et al*. 2020) and long-read assembly (11 Mb) (Lin *et al*. 2021). Plas-mid-copy number estimates based on Illumina and nanopore sequencing were in close agreement (**Figure 6D** and **Supplementary Figure 5**), validating our approach for estimating plasmid copy number and suggesting that the long-range sequencing assembly for CFJ157 is near-complete.

We determined structural features from the longest contigs and observed short tandem arrays (commonly 3-4 plasmids) interrupted by repetitive DNA sequences, presumably from the 1 kb ladder used as carrier DNA (Invitrogen) (**Figure 6B**). The sequence of the DNA ladder sequences is proprietary but are derived from phage DNA and *E. coli* DNA, including a ColE1 replication origin and an ampicillin resistance gene with homology to the injected plasmids. We observed structures that demonstrate array assembly by a combination of homologous recombination and NHEJ. In many cases, plasmid DNA was joined to ladder DNA sequences by homologous recombination in the amp or ori sequences. In another example, a tandem array of four plasmids (three *gfp* and one *tagRFP*) contained a breakpoint in the P*mlc-1* promoter. The two “parts” of the promoter were joined to the flanking ladder DNA with no obvious homology. We observed a similar structure where the tagRFP in a single plasmid (P*mlc-1*::tagRFP) was split into two fragments, consistent with prior results demonstrating that some plasmids reisolated from extra-chromosomal arrays contain truncations (Stinchcomb *et al*. 1985).

In conclusion, MosTI array insertion results in a diversity of fluorescence expression that is consistent with the plasmid-copy number in the integrated array. Integration does not generally cause wide-spread chromosomal aberrations (indels or duplications) near the insertion site, although some insertions influence the rescue marker, and possibly nearby genes, by position effect variegation. Arrays formed from a mix of supercoiled plasmids and linear carrier DNA contain short tandem plasmid arrays with some plasmids containing breakpoints in regulatory regions (promoter) or in coding regions.

## Discussion

We have described protocols and a set of reagents that enable <u>mo</u>dular <u>s</u>afe-harbor transgene insertion (MosTI) in the *C. elegans* genome. The insertion frequencies are high, and the isolation of transgenic strains with targeted insertions is facilitated by mutant rescue, antibiotic resistance, or fluorescence. MosTI is versatile by allowing insertion of both single-copy transgenes and extrachromosomal arrays. Finally, the reagents are modular with easy conversion between selection markers or generation of novel insertion sites.

### MosTI in comparison to commonly used insertion methods

In the past decade, several novel methods for transgene insertion have been described. We were inspired by these methods to develop MosTI that incorporate and improve on several of these advances. Advances have largely been driven by CRISPR/Cas9, with Dickinson *et al*. (2013) first showing that Cas9 induced double-strand DNA breaks could mediate single-copy transgene insertions at similar frequencies to MosSCI. Subsequently, improvements by many laboratories have increased the versatility and efficiency of CRISPR/Cas9 by, for example, using ribonucleoproteins (Paix *et al*. 2015; Au *et al*. 2019), oligos or modified repair templates (Dokshin *et al*. 2018), improved cloning strategies (Schwartz and Jorgensen 2016), and efficient proto-spacer sequences (Farboud and Meyer 2015). These improvements have, understandably, primarily focused on editing or tagging endogenous loci.

Some efforts have also focused on improving the ability to insert transgenes into safe-harbor locations. Aram *et al*. (2019) improved MosSCI by allowing the removal of selection markers. Silva-Garcia et al. (2019) used CRISPR/Cas9 to convert a set of MosSCI sites for use with *dpy-10* co-CRISPR editing to enrich for insertions (Arribere *et al*. 2014). This method, “SKI LODGE” is optimized for inserting PCR amplified coding regions with well-defined tissue specific promoters and protein tags present at the landing sites. The use of PCR products might be expected to allow multiplexed and high-throughput transgenesis but the low insertion efficiency (0.4 - 2%) of moderate size transgenes (2 kb) is currently a limitation. For this reason, we developed MosTI using plasmid templates but incorporated their use of designer landing sites with efficient protospacers.

Recently, Nonet (2020) demonstrated that recombination mediated cassette exchange (RMCE) using Flp/*FRT* or Cre/*LoxP* could be adapted for use in worms. In this case, landing sites with a GFP were randomly inserted into the genome by transposition (Frøkjær-Jensen *et al*. 2014). To insert a transgene by RMCE, the GFP is exchanged for a self-excising cassette (Dickinson *et al*. 2015) by screening for Rol animals. Although the use of RMCE is promising, particularly for inserting large transgenes, the method has a few limitations: First, insertions are generated at relatively low frequency (one in three injected animals). Second, co-insertion of the large self-excising cassette (7.7 kb) limits the size of transgene that can be inserted since most general cloning plasmids are limited to carrying 15 kb inserts (Bajpai 2014). Third, attempts by Nonet (2020) to integrate arrays were unsuccessful, possibly because constitutive expression of Flp recombinase “destroyed” arrays. Here, we have favored generating targeted landing sites by CRISPR/Cas9 and shorter selection markers. If preferred, the flexibility of modular MosTI landing sites can easily be converted to RMCE-compatible landing sites.

Finally, another research group has previously described a very similar use of a split antibiotic selection marker (HygroR) to insert single-copy transgenes (Stevenson *et al*. 2020). In this case, the authors generated a single landing site at the *ttTi5605* site on Chr. II and tested the efficiency of six protospacer sequences. Like our results, the authors note that single-copy insertions are easily identified based on hygromycin rescue although they report considerably lower insertion efficiencies (1 - 10%). Although insertion frequencies are difficult to compare across laboratories, one likely difference is our use of a recently developed germline-optimized Cas9 transgene (Aljohani *et al*. 2020). Stevenson *et al*. (2020) mainly focused on using *in vivo* recombineering of co-injected PCR fragments, which was first used for GFP tagging genes by Paix *et al*. (2016). *In vivo* recombineering has the significant advantage that cloning is minimized but comes at the expense of reduced insertion frequency and fidelity (Stevenson *et al*. 2020). We have preferred the use of clonal or synthetic transgenes but note that all MosTI landing sites and split selection markers are compatible with the *in vivo* recombineering strategies described by (Stevenson *et al*. 2020).

### Advantages of MosTI for single-copy insertions

MosTI has several advantages for single-copy insertions, in particular compared to MosSCI (Frøkjær-Jensen *et al*. 2008, 2012). First, the use of split selection markers makes it considerably easier to identify true transgene insertions, obviating the need for complex co-injection mixtures and time-consuming screens for insertions. Several negative selection markers have been developed to facilitate the identificaiton of single-copy insertion. The initial negative selection relied on killing array animals by expressing the toxin *peel-1* (Frøkjær-Jensen *et al*. 2012) but this method suffers from frequent escapers. A second method that relies on inducing paralysis by adding a Plas-mid to the injection mix which makes animals sensitive to histamine (El Mouridi *et al*. 2021) is less prone to escapers but requires the addition of drugs to every plate. In our own lab, we find that split selection is considerably easier, especially as it is often only necessary to generate a single extra-chromosomal array line with a heat-shock inducible Cas9, which makes it possible for even inexperienced injectors to generate several independent single-copy insertions with relative ease. For the same reason, it is difficult to compare the efficiency of MosSCI and MosTI; in our experience, most new students or researchers in the lab are quickly able to generate at least one extra-chromosomal array line (which enables both targeted single-copy and array insertion) whereas MosSCI insertions have typically been rather more difficult.

Each selection marker has its pros and cons. For *unc-19*, the selective pressure is very strong, it is easy to generate homozygous strains, and no specialized plates or fluorescence microscopes are required. The disadvantages include the difficulty of maintaining and injecting *unc-119* animals, no selection for the transgene in genetic crosses, and possible adverse effects of partial gene rescue. The advantages of antibiotic markers include easier strain handling, injection into wildtype animals, and strong selection in genetic crosses. The disadvantages include the cost and inconvenience of making plates with antibiotics and the difficulty of distinguishing homozygous from heterozygous inserts because non-rescued animals are “invisible” on the selective media. To our knowledge, fluorescent markers have not previously been used as the only selection marker for single-copy transgene insertions (in contrast to tagging endogenous genes), likely because fluorescence from arrays and insertions are difficult to distinguish. Split fluorescent selection markers are convenient because phenotypically normal animals are injected, homozygous and heterozygous inserts can be distinguished based on segregation, no special plates are needed, fluorescence protein expression is generally innocuous, and insertions are easily crossed to other strains by “following” the fluorescence. Minor limitations include the requirement for a fluorescence dissection microscope and possible interference between transgene and selection fluorophores. In this case, the fluorescent selection marker can be removed by Cre-mediated recombination.

Also, in contrast to MosSCI, the new MosTI insertion sites were selected based on chromosome location and local permissive chromatin marks. All the insertion sites show a high frequency of germline expression and consistently bright somatic expression. We have not attempted to do an in-depth phenotypic characterization of the strains but have not observed any obvious adverse phenotypes in animals with insertions. We have furthermore performed whole-genome sequencing on several of the insertion strains to detect mutations that were not out-crossed after the initial isolation of *unc-119*(*ed3*) or off-target effects from CRISPR/Cas9. We detected only a few mutations that cause amino-acid substitutions in genes with subtle phenotypes defined in **Supplementary Table 1**.

### Using MosTI for targeted array insertion

Targeted array integration by MosTI may be useful when high expression or co-expression of many plasmids is necessary. One striking example is the use of 41 different fluorescence drivers with overlapping expression that allow automated cell-specific identification and monitoring neuronal activity “NeuroPAL” (Yemini *et al*. 2021). To develop the NeuroPAL methodology, Yemini *et al*. (2021) tested many different plasmid combinations and generated integrated lines for many “landmark” fluorophores and a genetically encoded calcium sensor (GCaMP6). Each integrant required considerable effort, including integration, mapping, outcrossing, and phenotyping. Given the continuous development of improved fluorophores and optogenetic tools it is likely that tools such as NeuroPAL exemplify circumstances where rapid and reproducible array insertion would allow “prototyping” new functionalities (*e.g*., optogenetic inhibition or excitation of defined populations of cells). Similarly, somatic control of expression by Gal4 driver lines (Wang *et al*. 2017), FRT mediated gene activation (Davis *et al*. 2008; Voutev and Hubbard 2008), auxin-mediated degradation (Zhang *et al*. 2015), CRISPR inhibition (Long *et al*. 2015), or CRISPR activation (Gilbert *et al*. 2014) are being actively developed in *C. elegans* and often require sustained high transgene expression.

Our approach for inserting arrays using MosTI differ from the most commonly used methodology. Traditionally, arrays have been inserted by irradiation (Mello and Fire 1995; Evans 2006). Integrating arrays require specialized equipment, careful candidate screening, and is relatively slow. Furthermore, array insertion is random and inherently mutagenic. For these reasons, array integration is not standard practice despite mitotic stability and increased stability of expression in integrants (Evans 2006). In the past few years, two alternative array integration methods were developed. In one method, Noma and Jin (2018) adapted a genetically encoded reactive oxygen generator (“miniSOG”) to integrate arrays by light stimulation in just two weeks at high frequency (∼10%). Although this method is efficient, specialized equipment is still required, arrays are integrated randomly, and background mutations remain a concern. We have instead adapted a second method first developed by Yoshina *et al*. (2016) and recently expanded to additional gene loci (Yoshina and Mitani 2022) where arrays are integrated by CRISPR/Cas9. In this method, CRISPR/Cas9 is used to generate double-strand DNA breaks at a specific genomic location (*dpy-3* or *ben-1*) and in the array (typically, the AmpR marker). Arrays are integrated by non-homologous end-joining at relatively high frequencies (3 - 10%). The disadvantage is that arrays are inserted into coding regions causing a phenotype (Dpy) or using a somewhat complicated mix of positive (temperature selection with *vps-45*) and negative (benzimadole) leading to a relatively high frequency of false positives (Yoshina *et al*. 2016).

Here, we have developed a modified CRISPR/Cas9 based integration strategy that includes a positive selection marker which is compatible with the same safe-harbor locations used for MosTI insertions or with the *unc-19*(*ed3*) mutant strain. MosTI allows quick isolation (∼2 weeks) of integrated arrays at high frequency (7 to 40%) with no requirements for extensive screening or genetic mapping.

We characterized several array integrations using short-read sequencing and one array in detail by long-read Oxford Nanopore sequencing. From eight array integrations, we found no evidence for off-target insertions, large-scale chromosomal re-arrangements, or large indels at the integration sites, suggesting that targeted integrants are generally well-behaved. We observed unexpected effects in only one array integration (out of 39 in total), where the strain showed the hallmarks of position effect variegation of the *unc-119* rescue marker. Although we have not characterized this strain in detail, it is possible that repressive chromatin may partially silence *unc-119* at the junction between the genome and the integrant. To our knowledge, there are few published examples of somatic position effect variegation in *C. elegans*; targeted insertion of a large repetitive transgene structure could perhaps be used to understand heterochromatin spreading in worms that do not encode canonical genome organizers, such as CTCF (Heger *et al*. 2009).

Our long-read sequencing confirmed the tandem array structure and partial plasmid deletions in arrays, first demonstrated by Stinchcomb et al. (1985), and was in agreement with size estimates based on microscopy (Woglar *et al*. 2020) and sequencing a complex array (Lin *et al*. 2021). Although we have not sequenced arrays generated by linear DNA fragments, we and others have demonstrated better transgene expression in the soma (Etchberger and Hobert 2008) and germ line (Aljohani *et al*. 2020) from PCR products. Some of this difference may be driven by increased incorporation of linear fragments in arrays (Priyadarshini *et al*. 2022) but the presence of truncated promoters and fluorophores in the array also suggests that the cell may identify unusual DNA structures that are likely to lead to active silencing (Kelly *et al*. 1997; Leyva-Díaz *et al*. 2017). We envision that the ability to routinely insert arrays into specific locations will be useful for understanding basic characteristics of chromosomes, such as how chromosome size influences largescale genome domains (Liu *et al*. 2010), as well as enabling the initial steps in engineering synthetic *C. elegans* chromosomes.

### Multiplexed transgene insertion

CRISPR/Cas9 allows multiplexed transgene insertion because a single transgenic strain with an extra-chromosomal array can carry different repair templates. Kaymak *et al*. (2016) first demonstrated multiplexed repair (“library MosSCI”) to study the effect of 3’ UTRs, which largely determine the expression within the germline (Merritt *et al*. 2008). Similar to their observations, a complex mix of repair templates did not impair the MosTI repair process. However, library MosSCI was not inducible and required a relatively large number of injections to generate unique inserts (269 injected animals resulted in 11 unique transgene insertions) (Kaymak *et al*. 2016). Here, we show that coupling improved germline expression from arrays (Aljohani *et al*. 2020), an efficient sgRNA (Moreno-Mateos *et al*. 2015), and a split selection marker has allowed us to generate many insertions from a single injection by driving Cas9 expression using heat-shock. Our results were limited to proof-of-principle experiments using three plasmids that could be distinguished based on fluorescent protein expression. However, we propose that the method can be scaled up significantly: the rescuing part of the MosTI repair template (*cbr-unc-119* and the right homology region) is relatively short (<1 kb). Thus, a ∼5 Mb array can in principle carry approximately 2000 different copies of a short 1.5 kb transgene, more than enough room to encode a fluorophore with regulatory elements (*e.g*., promoters or 3’ UTRs). It should be possible to develop genetic screens that combine MosTI with large-scale gene synthesis, for example using oligo pools, and high-throughput sequencing to understand gene regulatory elements at the base-pair level (e.g., de Boer *et al*. 2020).

### MosTI curation and continuous development

Finally, standardized resources enable reproducibility across a scientific field, for example, by allowing direct comparison between different transgenes inserted at the same location or sharing of compatible reagents. Reagents for MosTI are freely available from the *Caenorhabditis* Genetics Center (CGC) and Addgene. To facilitate the uptake of transgenic methodology methods, we maintain a website (www.wormbuilder.org) with detailed descriptions of reagents and updated protocols. We designed MosTI to be highly modular and plan to continuously develop the technique, including new insertion sites, strains with multiplexed insertion sites each targeted by a different sgRNA, and new split selection markers. We hope that MosTI will be a resource that is of use to the many different types of experiments done by the *C. elegans* community.

## Acknowledgements

We thank Mohammed Aljohani for sharing unpublished reagents and Ramatoulaye Balde for excellent lab support. Research in C.F.-J.’s laboratory is supported by KAUST core funding and KAUST’s Office of Sponsored Research (OSR-CRG2020-4388). Some strains were provided by the CGC, which is funded by NIH Office of Research Infrastructure Programs (P40 OD010440).

## Contributions

Conceptualization: CFJ

Methodology: CFJ, SEM, FA

Software: FA

Formal analysis: SEM, FA

Investigation: SEM

Writing - original draft: CFJ

Writing - review & editing: CFJ, SEM, FA

Visualization: CFJ, SEM, FA

Supervision: CFJ

Funding acquisition: CFJ

## Data Availability

Short-read (Illumina) and long-read (Oxford Nanopore) sequencing data was deposited with Bio-Project number:

PRJNA827598

https://www.ncbi.nlm.nih.gov/sra

Full annotated plasmid sequences are included in

## Supplementary File 1

Plasmids are available from Addgene https://www.addgene.org/ and strains are available from the Caenorhabditis elegans Genetics Center (CGC) https://cgc.umn.edu/

## Materials & Methods

### Strains

Strains were maintained using standard methods (Brenner 1974) and were grown at 20°C on OP50 or HB101 bacteria. Strains containing MosTI landing sites have been deposited with the *Caenorhabditis* Genetics Center (CGC) (see **Supplementary Table 1**).

### Commercial software

Some figures were created with BioRender.com. Figures were designed using Adobe Illustrator 2022 (v26.0.2). Statistical analysis was performed using GraphPad Prism 9 for macOS (v9.3.1). The manuscript was written in Microsoft Word for Mac (v.16.60) and tables were generated using Microsoft Excel for Mac (v. 16.60). In silico molecular biology was performed using ApE (A plasmid Editor)(Davis and Jorgensen 2022) which is freely available at https://jorgensen.biology.utah.edu/wayned/ape/

### Molecular biology

Plasmids were generated by standard molecular techniques, including three-fragment multisite Gateway reactions (Invitrogen cat. no. 12538200), Gibson assembly (Gibson *et al*. 2009), Golden-Gate cloning (Engler *et al*. 2009), or by gene synthesis (Twist Bioscience, CA, USA). All PCRs were performed using a high-fidelity DNA polymerase (Phusion, New England Biolabs, F530S) and constructs generated by PCR were sequence verified by Sanger sequencing. Most plasmids have been deposited with Addgene (Cambridge, MA, USA) (see **Supplementary Table 1**). Annotated GenBank files of all plasmids are included in **Supplementary File 1**.

### MosTI landing sites

Please see **Supplementary Table 1** for the exact insertion sites and sgRNA sequences.

### *unc-119* selection

We generated target plasmids containing a codon-optimized synthetic *unc-119* rescue marker (“*syn-unc-19(*+)”) flanked by 250 bp of homology to each side of the insertion site (pSEM203 Chr. II, pSEM224 Chr. I, pSEM226 Chr. IV). We injected these targeting plasmids (individually) into *unc-119*(*ed3*) animals at 25 ng/ul with 25 ng/ul pCFJ2474 (P*smu-2*::Cas9::*gpd-2*::*mCherry*), 15 ng/ul of sgRNA (specific to each site) and 10 ng/ul of fluorescent co-injection markers pSEM233 (P*mlc-1*::*tagRFP*) or pSEM231 (P*mlc-1*::*gfp*) (El Mouridi *et al*. 2020) for a final DNA concentration of 100 ng/ul. After injection, P0 plates were placed at 25°C to increase Cas9 expression. The screen for Unc rescue was done after starvation. Strains that were homozygous for the landing sites were subsequently injected with a plasmid encoding Cre recombinase (pMDJ39) to remove the synthetic *unc-119* rescue marker (flanked by LoxP sites) and identified by screening for Unc animals. We validated all MosTI insertions by PCR and Sanger sequencing. Additionally, we whole-genome sequenced two MosTI insertion strains, CFJ77 (chr. I) and CFJ42 (chr. II) to identify background mutations (see **Supplementary Table S1**). All other insertions strains (for example, strains with split hygroR and P*mlc-2*::*gfp* selection) were derived from the same genetic background (originally an *unc-119*(*ed3*) mutant strain) and are expected to share the same background SNPs.

### Converting MosTI landing site to a different selection marker

We converted MosTI strains with split *unc-119* selection to split P*mlc-2*::*gfp* and hygromycin selections. MosTI strains for a given chromosomal location with *unc-119* selection were injected with a conversion plasmid (pSEM258 for hygroR and pSEM260 for P*mlc-2*::*gfp*) and standard plasmids used for MosTI single-copy insertions (Cas9, *hygroR*, sgRNA, *tagRFP* or *gfp* fluorescent co-injection markers). After injection, P0 animals were placed at 25°C. After 48 hours, 500 ul of a 4 mg/ml hygromycin solution was added to the NGM plates to select for transgenic animals. Single-copy insertion of the conversion transgenes was identified by *unc-119* rescue after the food was exhausted. All converted MosTI sites were validated by Sanger sequencing. The conversion strategy should, in principle, work for any selection marker that can be split into two non-rescuing fragments.

### Single-copy transgene insertion using MosTI

#### Constitutive Cas9 protocol

In this protocol, a codon-optimized Cas9 containing PATCs in introns to prevent silencing and a fluorophore to monitor expression (pCFJ2474, P*smu-2*::*cas9*::*gpd-2 tagRFP-T*) (Aljohani *et al*. 2020) was used for constitutive expression from extrachromosomal arrays. The injection mix consisted of 25 ng/ul linearized Cas9 (pCFJ2474), 25 ng/ul target vector (various), 15 ng/ul pCFJ782 (HygroR), 15 ng/ul sgRNA (pSEM318), and 10 ng/ul of fluorescent co-injection markers (P*mlc-1*::*tagRFP*, pSEM233 or P*mlc-1*::*gfp*, pSEM231) for a total DNA concentration of 100 ng/ul. The plasmids were injected into young adult Unc animals grown at 15°C or 20°C from strains with MosTI landing sites. Single, injected animals were placed on NGM plates seeded with OP50 and incubated at 25°C. 48 hours after injection, 500 ul of a 4 mg/ml hygromycin solution was added to P0 plates to select for transgenic animals. Single-copy insertions were identified by screening plates for any moving animals (*unc-119* rescue) once the food was exhausted (approximately seven to ten after injection).

#### Heat-shock protocol

In this protocol, codon-optimized Cas9 was expressed under a heat-shock promoter for inducible Cas9 expression. The injection mix contained 25 ng/ul of P*hsp*-*16.41*::Cas9 (pMDJ231) linearized with ApaLI, 25 ng/ul target vector, 15 ng/ul pCFJ782 (HygroR), 15 ng/ul sgRNA (pSEM318), and 10 ng/ul of muscle co-injection markers (P*mlc-1*::*tagRFP*, pSEM233 or P*mlc-1*::*gfp*, pSEM231 for a total DNA concentration of 100 ng/ul. The plasmids were injected into young adult Unc animals grown at 15°C or 20°C from strains with MosTI landing sites. Single, injected animals were placed on NGM plates seeded with OP50 and incubated at 25°C. 48 hours after injection, 500 ul of a 4 mg/ml hygromycin solution was added to P0 plates to select for transgenic animals. Two different heat-shock protocols were used to generate single-copy insertions: (a) 37°C for 1 hour or (b) 30°C for 18 hours (Nonet 2020). Animals with single-copy insertions were identified based on *unc-119* rescue after the food was exhausted. Note, we frequently observed transgene insertions prior to heat-shock, suggesting that the heat-shock promoter is leaky or that transgenic animals are under stress (*e.g*., due to temperature, antibiotic selection, or starvation). The longer heat-shock protocol (30°C for 18 hours) was slightly more efficient and was more convenient as all plates placed in an incubator will be equilibrated to the surrounding temperature without the need for wrapping plates in parafilm and using a water bath (Boulin & Bessereau, 2007) or splitting plates into a single layer (Frøkjær-Jensen et al., 2014).

#### Heat-shock induced MosTI multiplex insertions over generations

A MosTI strain containing a landing site on Chr. II (CFJ42) was injected with a mix containing 10 ng/ul linearized transgene(s), 10 ng/ul pCFJ782 (HygroR), 10 ng/ul pSEM318 (sgRNA), 10 ng/ul pMDJ231 (P*hsp*-*16.41*::Cas9) linearized with ApaLI, and DNA ladder (1 kb ladder, Thermo Fisher Scientific, cat. no. SM1331) added to a final concentration of 100 ng/ul. Injected animals were singled to NGM plates seeded with OP50 and placed at 25°C. One day after injection, 500 ul of 4 mg/ml hygromycin was added to injection plates to select for transgenic strains. The experiment was performed in parallel with seven independent transgenic strains. For each transgenic strain, a single non-rescued transgenic Unc animal was picked to an NGM plate seeded with OP50 seven days after injection and placed at 25°C. For every generation, a single non-rescued transgenic animal was transferred to a new NGM plate with OP50 before the food was exhausted. Subsequently, the transgenic animals were heat-shocked to induce Cas9 expression seven days after being singled to NGM plates. The plates were screened for single-copy transgene insertions (based on moving *unc-119*-rescued animals) after the food was exhausted. In some experiments, the injection mix contained several target transgenes (P*rpl-7A*::*gfp*, P*eft-3*::*mMaple*, and P*eft-3*::*mScarlet*). We determined which transgene had been inserted based on fluorescence, PCR, and Sanger sequencing.

### Extrachromosomal array insertions

Extra-chromosomal arrays can be inserted into MosTI sites or into the endogenous *unc-119* locus (in *ed3* mutants) using *unc-119* selection. We have also generated targeting fragments to insert arrays into MosTI sites that use hygromycin and P*mlc-2*::*gfp* but have not tested these fragments (although we deposited the reagents with Addgene). The protocols are identical except for using targeting fragments and sgRNAs that are specific to each insertion site (listed in **Supplementary Table 1**). Here we describe validated array insertion protocols based on *unc-119* rescue.

#### One-step direct plasmid-based array integration

With this strategy, all injected plasmids are “directly” integrated at the target site, including plasmids expressing Cas9 and the sgRNA. We injected adult *unc-119* hermaphrodites with a mix composed of 10 ng/ul of pCFJ2474 (P*smu-2*::*cas9*::*gpd-2 tagRFP-T*), 5 ng/ul of array target fragment (*e.g*., pSEM371 for integration at the endogenous *ce-unc-119* locus), 15 ng/ul of sgRNA targeting the integration site and the array target fragment (*e.g*., pSEM376 for *ce-unc-119*), 10 ng/ul of transgenes for integration, and DNA ladder (1 kb plus, Invitrogen) to a final concentration of 100 ng/ul. Injected animals were placed on NGM plates and grown at 25°C degrees. Seven to ten days after injection, we screened plates for array integration based on *unc-119* rescue and expression of injected transgenes (typically, fluorescent markers). We treated any plate with rescued animals as a single independent integration event and picked only a single clonal strain derived from any injected animal.

#### One-step plasmid-based array integration in pre-injected strain

With this strategy, only the transgenes, the array target fragment, and the sgRNA plasmid were integrated. First, a MosTI strain containing a landing site on Chr. II (CFJ42) was injected with a mix containing: 25 ng/ul pMDJ231 (P*hsp*-*16.41*::Cas9) linearized with ApaLI, 15 ng/ul pCFJ594 (NeoR), 10 ng/ul pSEM235 (P*mlc-1*::*mCherry*), and DNA ladder (1 kb ladder, Thermo Fisher Scientific) added to a final concentration of 100 ng/ul. Injected animals were singled to NGM plates seeded with OP50 and placed at 25°C. One day after injection, 500 ul of 25 mg/ml neomycin was added to injection plates to select for transgenic strains (not phenotypically rescued for *unc-119*(*ed3*)). A single strain was selected as an injection strain for a second step, where adult animals were injected with a mix containing: 15 ng/ul pSEM376 (sgRNA), 15 ng/ul pCFJ782 (hygroR), 15 ng/ul pSEM234 (P*mlc-1::mCherry::NSL*), 15 ng/ul pSEM233 (P*mlc-1::gfp*), 5 ng/ul pSEM371 (integration fragment), and 1 kb DNA ladder (Invitrogen) to a total concentration of 100 ng/ul. Single, injected animals were placed on NGM plates seeded with OP50 and incubated at 25°C. 48 hours after injection, 500 ul of a 4 mg/ml hygromycin solution was added to P0 plates to select for transgenic animals. Seven to ten days after injection (before the bacterial lawn was exhausted), strains with the two extra-chromosomal arrays were heat-shocked (30°C for 18 hours) and screened for *unc-119* rescue one generation later. We subsequently allowed the initial array (containing NeoR and P*mlc-1*::*mCherry*) to be lost on standard NGM plates with no antibiotic selection and the integrants were rendered homozygous. Cas9 should only cut the array formed in the second injection and as expected, we only observed antibiotic resistance and markers from the second injection mix in the integrated strains.

#### Two-step plasmid-based array integration

With this strategy, only the transgenes and the array target fragment are integrated; plasmids encoding Cas9 and the sgRNA are injected in a second round and are not part of the integrated array. As a first step, we generated transgenic lines by injecting a mix composed of 2.5 ng/ul of the array target fragment (*e.g*., pSEM371 for *ceunc-119* integration), 15 ng/ul of pCFJ782 (HygroR), 10 ng/ul of transgenes, and DNA ladder (1 kb plus, Invitrogen) to a final concentration of 100 ng/ul. Injected animals were singled to NGM plates seeded with OP50 and placed at 25°C. One day after injection, 500 ul of 4 mg/ml hygromycin was added to injection plates to select for stable transgenic strains. In the F2 or F3 generation we picked a single clonal animal to a seeded NGM plate with hygromycin and expanded the population (note that these animals are not rescued for the *unc-119* phenotype but are hygromycin resistant). For the second step, we injected Unc transgenic animals carrying extra-chromosomal arrays with a mix containing 20 ng/ul pCFJ2474 (P*smu-2*::*cas9*::*gpd-2 tagRFP-T*), 15 ng/ul of plasmid expressing an sgRNA to cut the integration site and the array fragment (*e.g*., pSEM376 for *ce-unc-119* integration), 10 ng/ul of fluorescent co-injection marker (pSEM233 or pSEM231), and DNA ladder (1 kb plus, Invitrogen) to a final concentration of 100 ng/ul. After injection, we placed single individual injected animals on seeded NGM plates with hygromycin and grew the animals at 25°C. We identified strains with array integrations seven to ten days after injection based on unc-119 rescue. We treated any plate with rescued animals as a single independent integration event and picked only a single clonal strain derived from any injected animal.

#### Two-step protein-based array integration

With this strategy, only the transgenes and the array target fragment are integrated; Cas9 protein (IDT) and crRNA/tracrRNA (IDT) are injected in a second round. Integrations are generated faster with this protocol compared to plasmid-based protocols as array integrants are identified in the progeny of the injected animals. As for the two-step plasmid protocol, we first generated transgenic lines by injecting a mix composed of 2.5 ng/ul of the array target fragment (*e.g*., pSEM371 for *ce-unc-119* integration), 15 ng/ul of pCFJ782 (HygroR), 10 ng/ul of transgenes, and DNA ladder (1 kb plus, Invitrogen) to a final concentration of 100 ng/ul. Injected animals were singled to NGM plates seeded with OP50 and placed at 25°C. One day after injection, 500 ul of 4 mg/ml hygromycin was added to injection plates to select for stable transgenic strains. In the F2 or F3 generation we picked a single clonal animal to a seeded NGM plate with hygromycin and expanded the population (note that these animals are not rescued for the *unc-119* phenotype but are hygromycin resistant). For the second step, we prepared guide RNA duplexes *in vitro* by incubating 3 ul of 100 uM crRNA (*e.g*., targeting *ce-unc-119*), 3 ul 100 uM tracrRNA, and 4 ul nuclease-free duplex buffer (IDT) at 95°C for 5 min, followed by a 5 min incubation at room temperature. We mixed 3 ul of the crRNA::tracrRNA duplex with 10 ug recombinant Cas9 protein (Alt-R S.p. Cas9 Nuclease V3, IDT), 10 ng/ul fluorescent co-injection marker (pSEM233 or pSEM231), and nuclease-free water to a final volume of 10 ul. We injected transgenic Unc animals carrying extra-chromosomal arrays with this Cas9 protein mix and placed individual injected animals on NGM plates seeded with OP50 at 25°C. Rescued animals were identified in the progeny of the injected animals (the F1 generation) and isolated before the food was exhausted. We treated any plate with rescued animals as a single independent integration event and picked only a single clonal strain derived from any injected animal.

### Characterization of arrays integrated by MosTI

#### Quantification of fluorescence expression using COPAS large-particle flow cytometer

We obtained similar mixed populations by placing four L4 worms on small NGM plates seeded with OP50 and used four plates per strains. The animals were grown at 25°C for seven days and then washed off with a M9 buffer and placed on ice. After 10 min, the pellet was washed with water to remove bacteria. We quantified the fluorescence of each strain using a COPAS flow cytometer (Union Biometrica). Between each strain, we washed all the tubing by running water for at least one minute. Settings for one-step insertion: Gain = 2, green: 400 volts; red: 450 volts. Settings for two-step insertion: Gain = 2, green: 300 volts; red: 400 volts. Both green and red fluorescence were expressed along the entire length of the worm, so we based our quantification on the integral of the fluorescence signal. We selected animals with a TOF between 600 and 900 (mainly young adults) and with an extinction coefficient below 35000 to select for animals that came straight through the flow cell. We manipulated the data in Microsoft Excel (version 16.59) and graphed the data with GraphPad Prism (version 9.3).

#### Background SNP analysis in injection strains

Raw sequencing reads were trimmed to remove adapters and filtered for low quality bases (Fastqc) (Andrews 2010). The filtered reads were aligned to the C. *elegans* reference genome (WBcel235) with BWA V0.7.17 (Li 2013). Single-nucleotide variants and indels were called with GATK4 Haplotypecaller (Poplin *et al*. 2018) using filters: QD < 2.0, FS > 60, MQ < 40, SOR > 4, MQRankSum < −8, ReadPosRankSum < −8. Next, we used snpEff to filter out SNPs and indels outside proteincoding genes and those with “modifier” impact (Cingolani *et al*. 2012).

#### Plasmid copy-number estimation

Genomic DNA from eight strains with integrated arrays was isolated using a DNeasy blood and tissue kit according to the manufacturer’s instructions (Qiagen, cat. no. 69504). Whole-genome sequencing (150 bp paired end) was performed by Novogene. Short Illumina reads were mapped to the unique regions of integrated transgenes with BWA V0.7.17 (Li 2013). We calculated copy-number estimates with the formula: (mapped transgene reads * genome size) / (total genomic mapped reads / mapped transgene length). All copy-number estimates are per haploid genome.

#### Integrated array assembly from Oxford Nanopore sequencing

High molecular weight genomic DNA was extracted from CFJ157 (*kstIs6* [pSEM371; pSEM233; pSEM231, 1kb ladder] III) using a MagAttract HMW DNA kit following the manufacturer’s instructions (Qiagen, cat #67563). Oxford Nanopore sequencing was performed by Novogene (China) using a PromethION instrument. We removed *C. elegans* reads by mapping base-called and quality filtered reads to the *C. elegans* reference genome (WBcel235) with transgene sequence homology and 40 kb of the reference sequence flanking the insertion site masked. Unmapped reads (corresponding to the integrated array, flanking genomic sequences and *E. coli* genome) were assembled with the Canu long-read assembler using default settings except for: ‘corOutCoverage=9999’ ‘minOverlapLength=2500’ ‘minReadLength=2500’ ‘corMinCoverage=0’ ‘corMaxEvidenceErate=0.15’ (Koren *et al*. 2017). Array contigs were identified by aligning integrated transgene sequences to contigs with MUMmer V4 (Marçais *et al*. 2018). A single contig (∼5Mb) corresponding to the *E. coli* genome was excluded from further analysis.

## Supplementary Materials

**Supplementary Figure 1.**
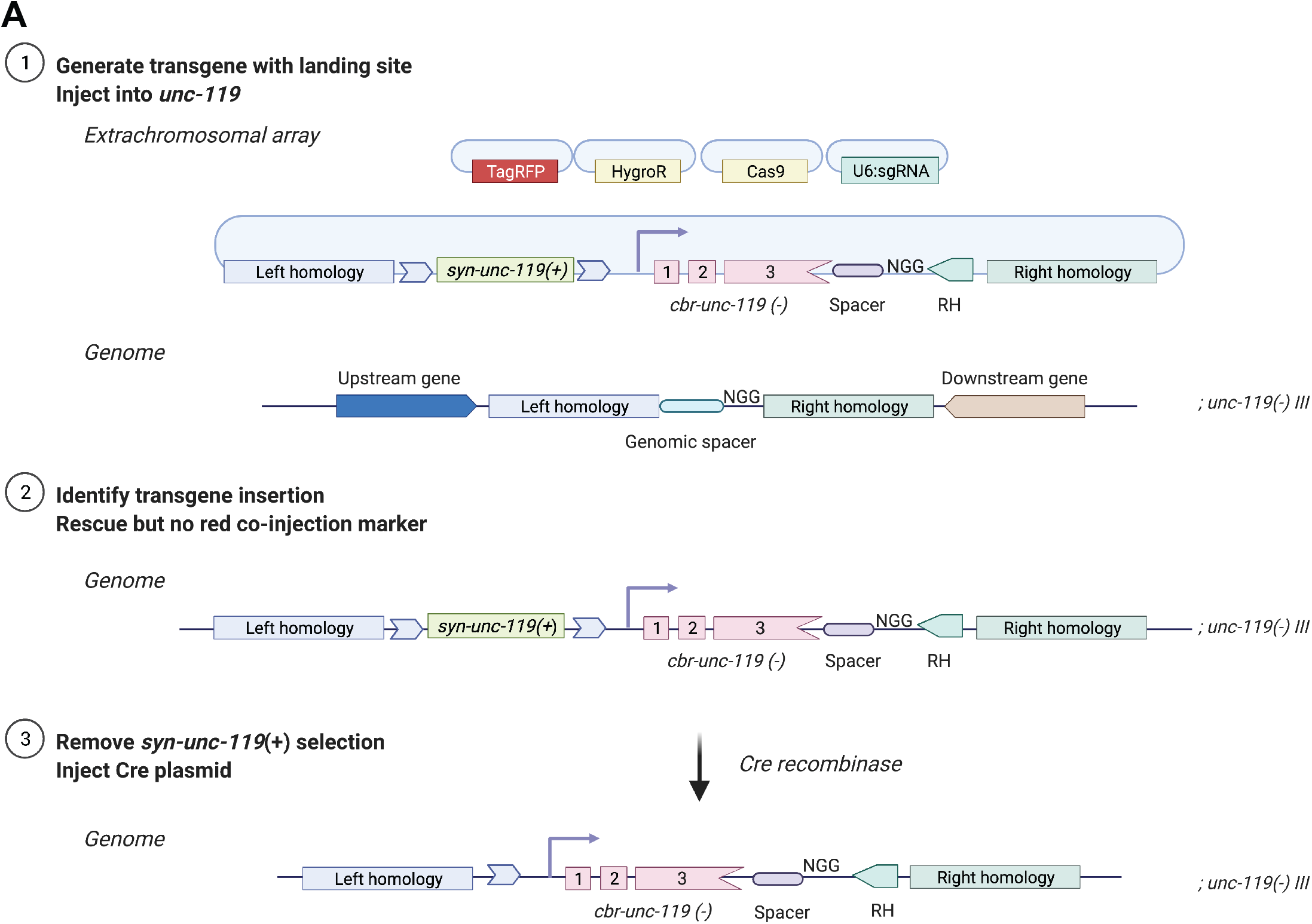
| Two-step protocol to generate initial MosTI landing sites **A**. 1. As a first step, we generated a targeting construct that contains homology regions (left and right homology) to the insertion site, a coding-optimized *unc-119* rescue fragment (“*syn-unc-119(+)*”), the split *cbr-unc-119* marker (“*cbr-unc-119(-)*”), a spacer sequence, and a right homology region for transgene insertions (“RH”). We selected sites in permissive chromatin domains (Ho *et al*. 2014) and between two convergently transcribed genes to minimize the effect of nearby promoters and enhancers. The targeting construct, fluorescent co-injection markers (“tag-RFP”), an antibiotic selection marker (“HygroR”), plasmids expressing Cas9, and sgRNA are injected into a strain with the *unc-119*(*ed3*) mutation. 2. Animals with the landing site inserted at the target genomic location were identified based on Unc rescue and loss of the extra-chromosomal array (based on loss of fluorescent co-injection markers). 3. Finally, *syn-unc-119(+)* selection was removed from the landing site by injecting a plasmid that expresses Cre recombinase and screening for Unc animals.

**Supplementary Figure 2.**
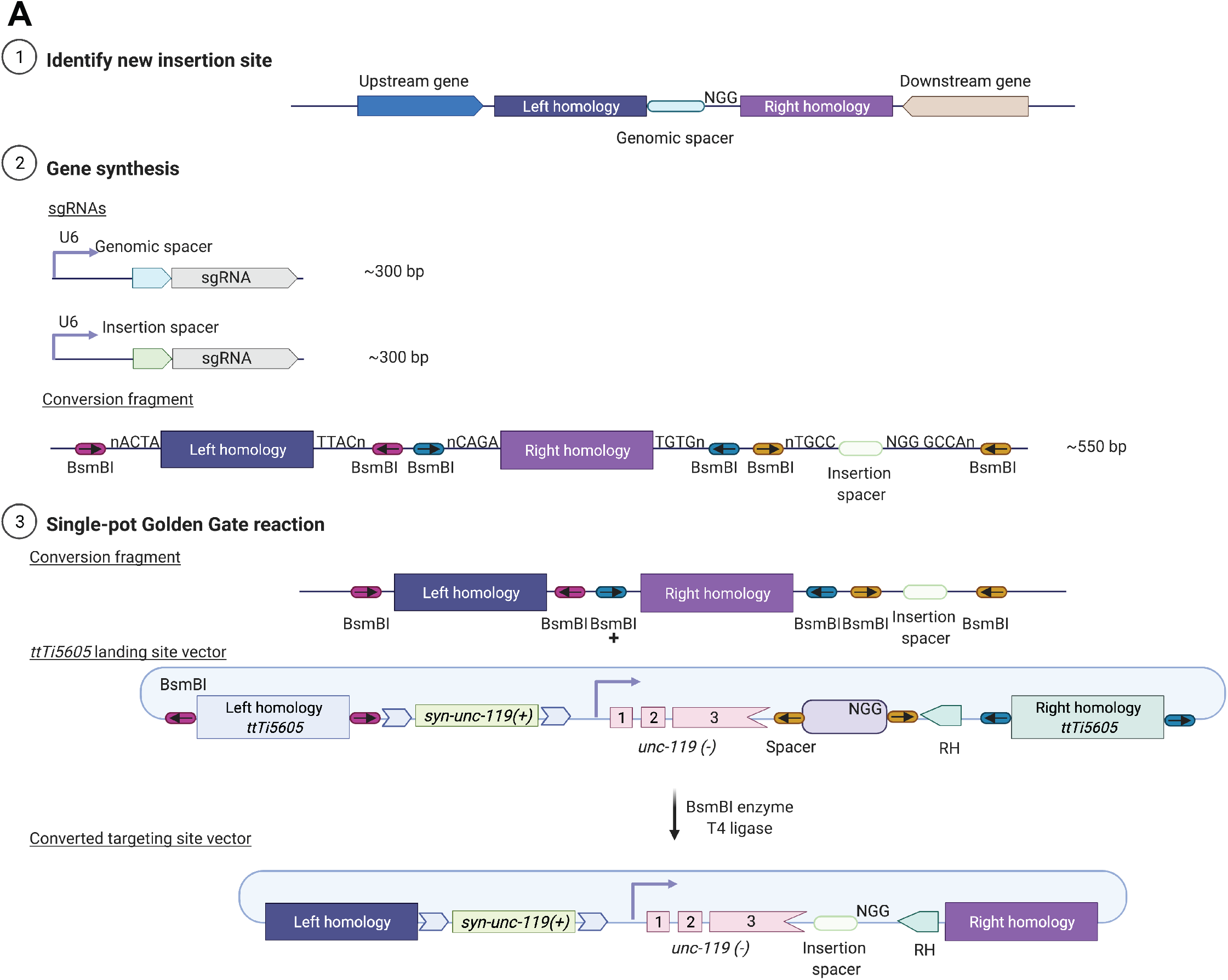
| Generating new MosTI landing site. **A**. 1. As a first step, we identified appropriate insertion sites. For “safe-harbor” landing sites, we selected insertion sites located in permissive chromatin (Ho *et al*. 2014) and between the 3’UTRs of two convergently transcribed endogenous genes. 2. In a second step, we synthesized two sgRNAs: one for inserting the landing site and one for inserting transgenes at the landing site (optional - not required if using the same sgRNA as used for the standard set of MosTI landing sites). We synthesized a conversion fragment containing left and right homology regions (250 - 500 bp) and a novel insertion spacer flanked by the appropriate Golden-Gate cloning overhangs (indicated in figure). Note: any BsmBI sites in the homology regions and spacer must be removed. 3. We performed a Golden-Gate reaction using BsmBI between the conversion fragment and the *ttTi5605* MosTI landing site vector (pSEM203). This reaction exchanged the homology arms and the insertion site spacer but maintained the codon-optimized *syn*-*unc-119(+)* and split *cbr-unc-119* selection marker.

**Supplementary Figure 3.**
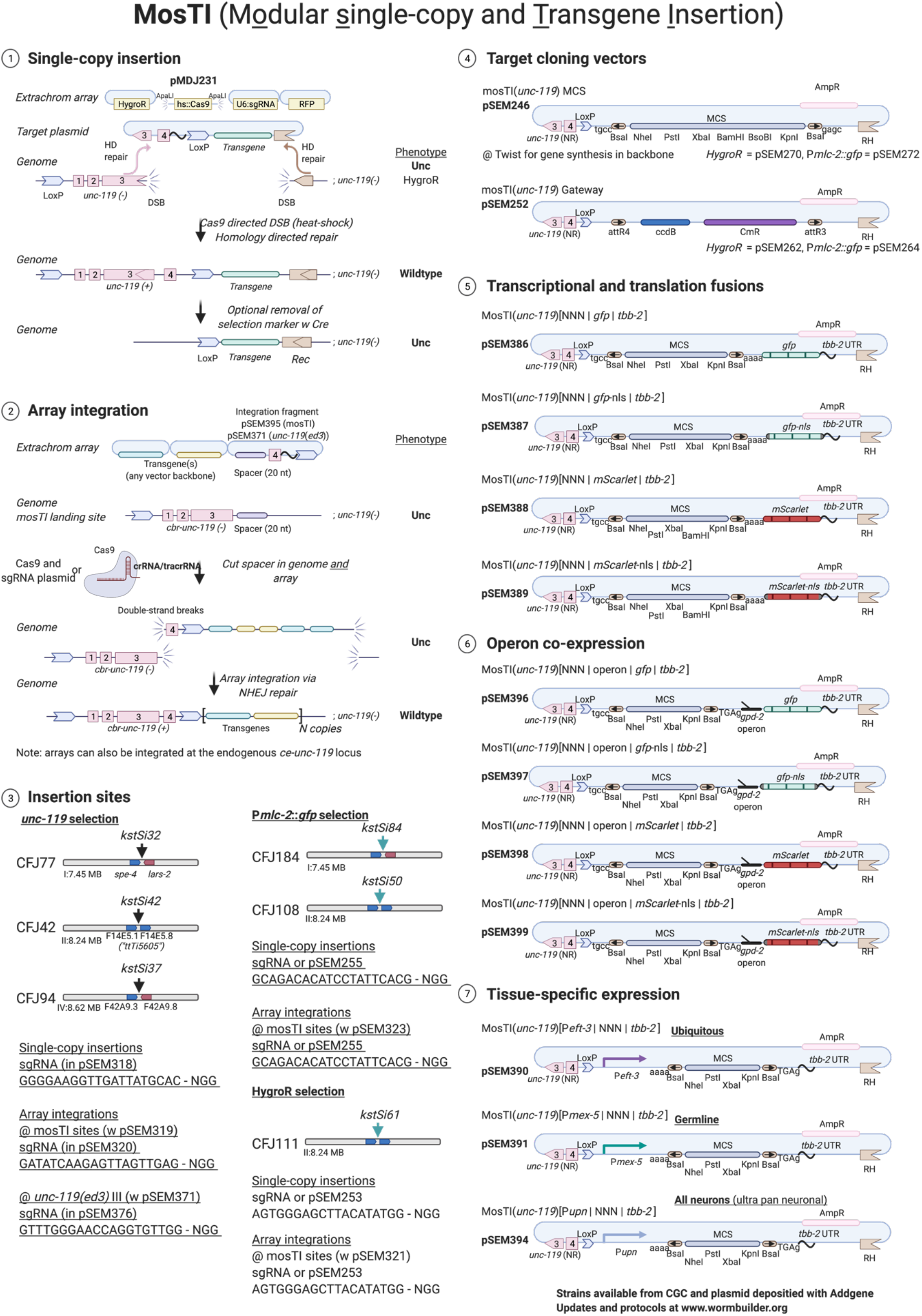
| MosTI overview. This figure contains a single overview of MosTI methods and reagents. 1. Schematic overview for generating MosTI single-copy insertions. 2. Schematic overview for generating MosTI array integrations. 3. Overview of single-copy MosTi sites. 4. Cloning vectors for MosTi single-copy insertions. All vectors are compatible with restriction enzyme cloning into the multiple cloning site (MCS) or Golden-gate based cloning using BsaI. Note that pSEM246 has been deposited with Twist Bioscience for direct gene synthesis. 5. MosTI vectors for transcriptional and translation fusions (*unc-119*-based selection). 6. MosTI vectors for fluorophore co-expression using the *gpd-2* operon. Transgene expression can be monitored by fluorophore co-expression. 7. Tissue-specific MosTI expression vectors. P*eft-3* is ubiquitously expressed (including the germline), P*mex-5* is germline specific (Zeiser et al. 2011), and Pupn is pan-neuronal (“ultra pan neuronal”)(Yemini et al. 2021).

**Supplementary Figure 4.**
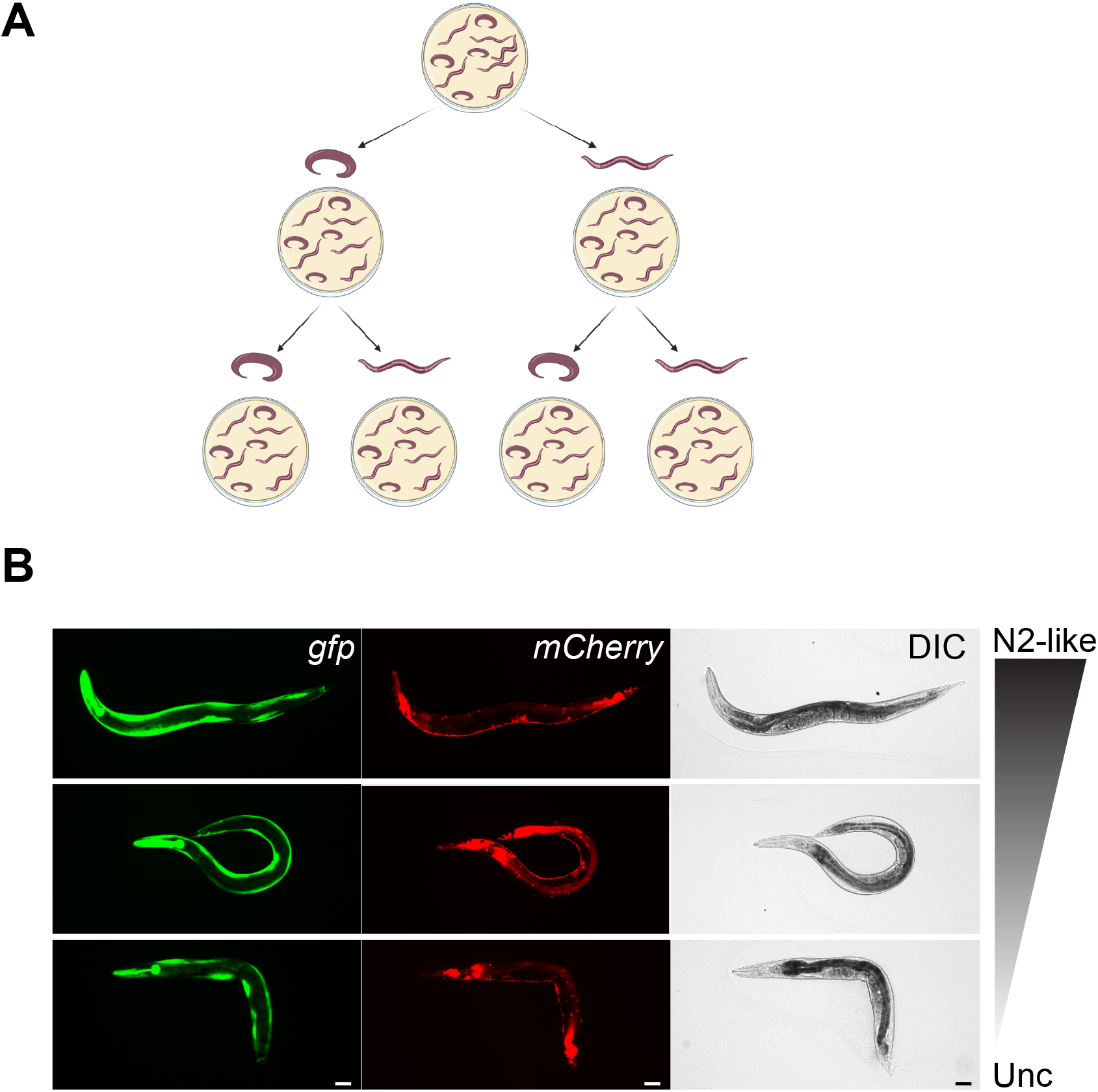
| Position effect variegation following array integration. **A**. Schematic overview of position effect variegation after targeted integration of an extra-chromosomal array into a MosTI site by the “two-step heat-shock plasmid” protocol. We observed *unc-119* rescue indicating that an NHEJ event had happened between the genomic location and the array, but we were unable to generate a homozygous rescued strain. We observed a continuum of phenotypes from fully rescued to non-rescued animals suggesting phenotypic variegation. Non-rescued animals would segregate a mix of rescued and non-rescued animals, like the segregation pattern of rescued animals. **B**. Fluorescence images of rescued and non-rescued transgenic animals. All animals contained the injected fluorescence markers regardless of the degree of rescue. We were also unable to detect any obvious mosaicism suggesting that the strains did not contain an extra-chromosomal array with unusual segregation. Scale bar = 20 microns, 10x magnification.

**Supplementary Figure 5.**
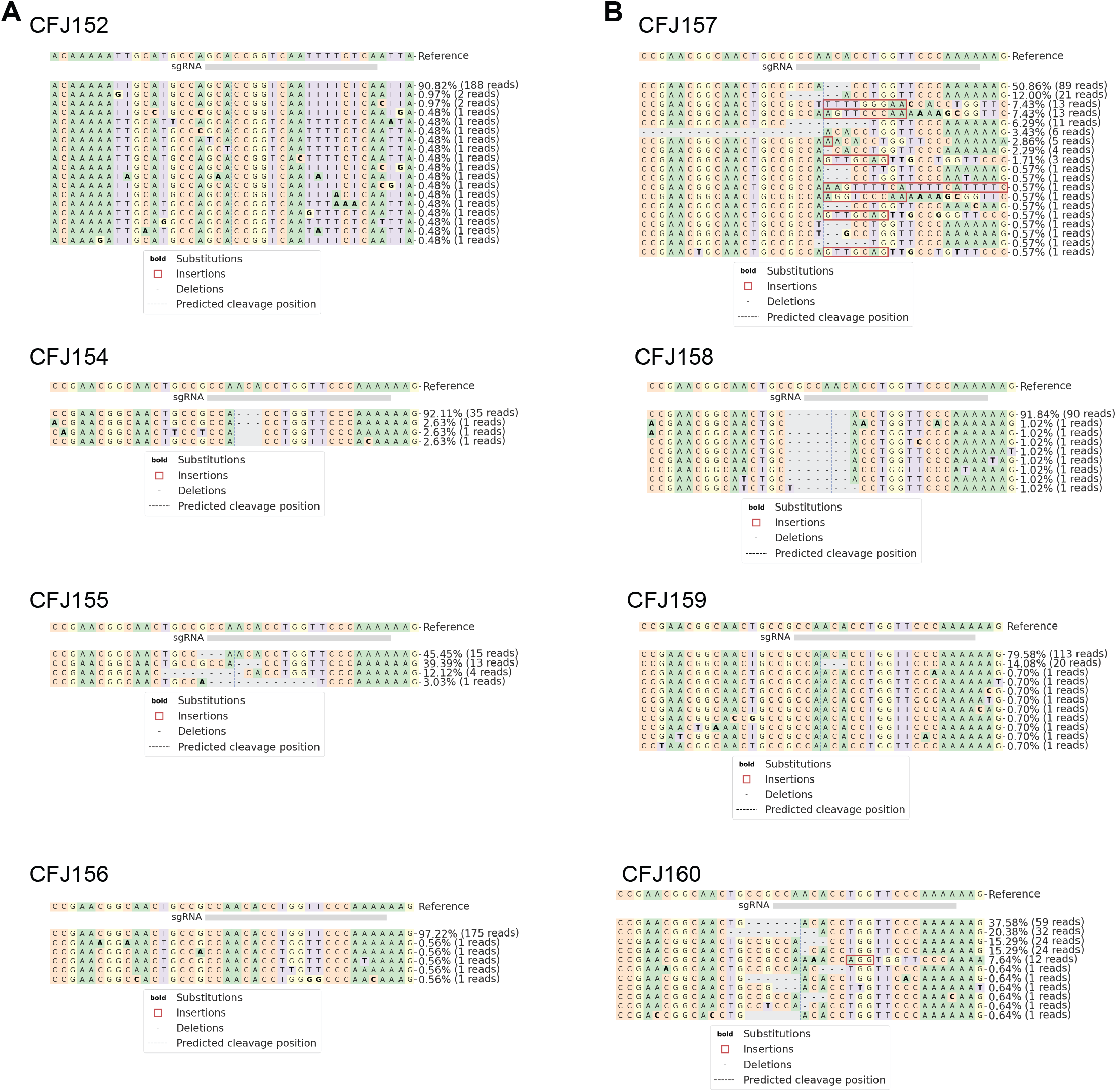
| Indels in integration fragments within integrated arrays. We analyzed Illumina short-read whole-genome sequence data (150 bp paired-end reads) using CRISPResso2 (Clement *et al*. 2019) to determine the integrity and copy-number number of the integration fragment within MosTI integrants. The integration fragments (and the genomic location) are cut by the sgRNA, and parts of the extra-chromosomal array are integrated by non-homologous end-joining. The number of different alleles with reasonable read coverage (>1 read) are a reasonable lower bound for the copy-number of the integration plasmid. See **Figure 6A** for plasmid-copy number estimates based on sequence coverage. **A**. Array integrations generated by the “one-step plasmid” protocol. **B**. Array integrations generated by the “two-step protein” protocol.

**Supplementary Figure 6.**
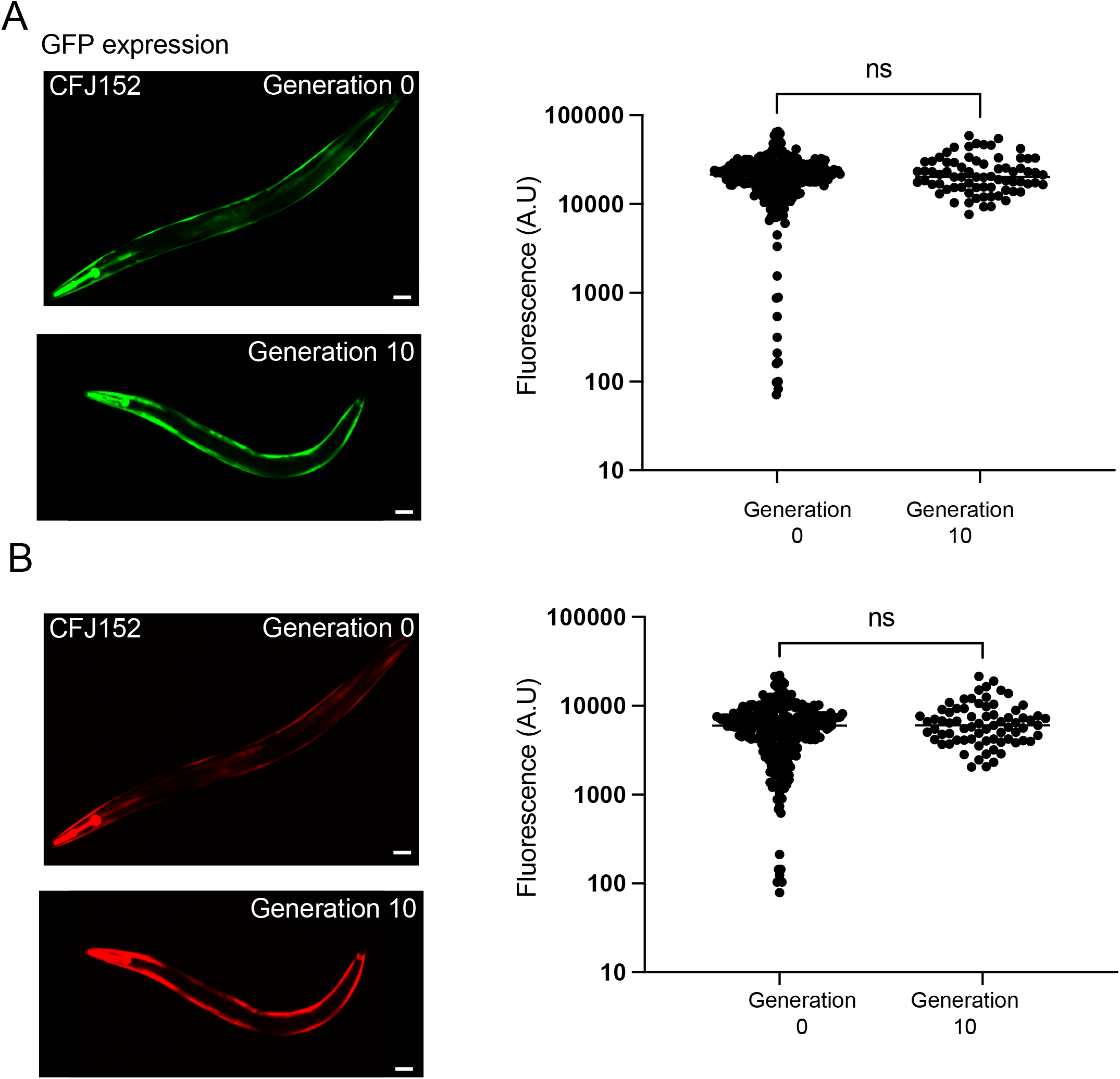
| Fluorophore expression is unchanged after ten generations for integrated array. We propagated the strain CFJ152 (*kstIs1*[pCFJ2474 (Cas9); pSEM371(integration fragment); pSEM376 (sgRNA1), pSEM377(sgRNA2);pSEM233 (P*mlc-1*::*tagRFP*); pSEM231 (P*mlc-1*::*gfp*), 1kb ladder] III) for ten generations without selecting for any fluorescence to determine if the integrated array changed over time. **A**. GFP expression at generation 0 and generation 10. **B**. TagRFP expression at generation 0 and generation 10. Left, fluorescence images of adult animals. Scale bar = 20 microns. Right. Quantification of fluorescence levels using COPAS large-particle flow sorter (animals selected with time-of-flight between 700-900 to select mainly adult animals). Statistics: Mann-Whitney t-test.

**Supplementary Figure 7.**
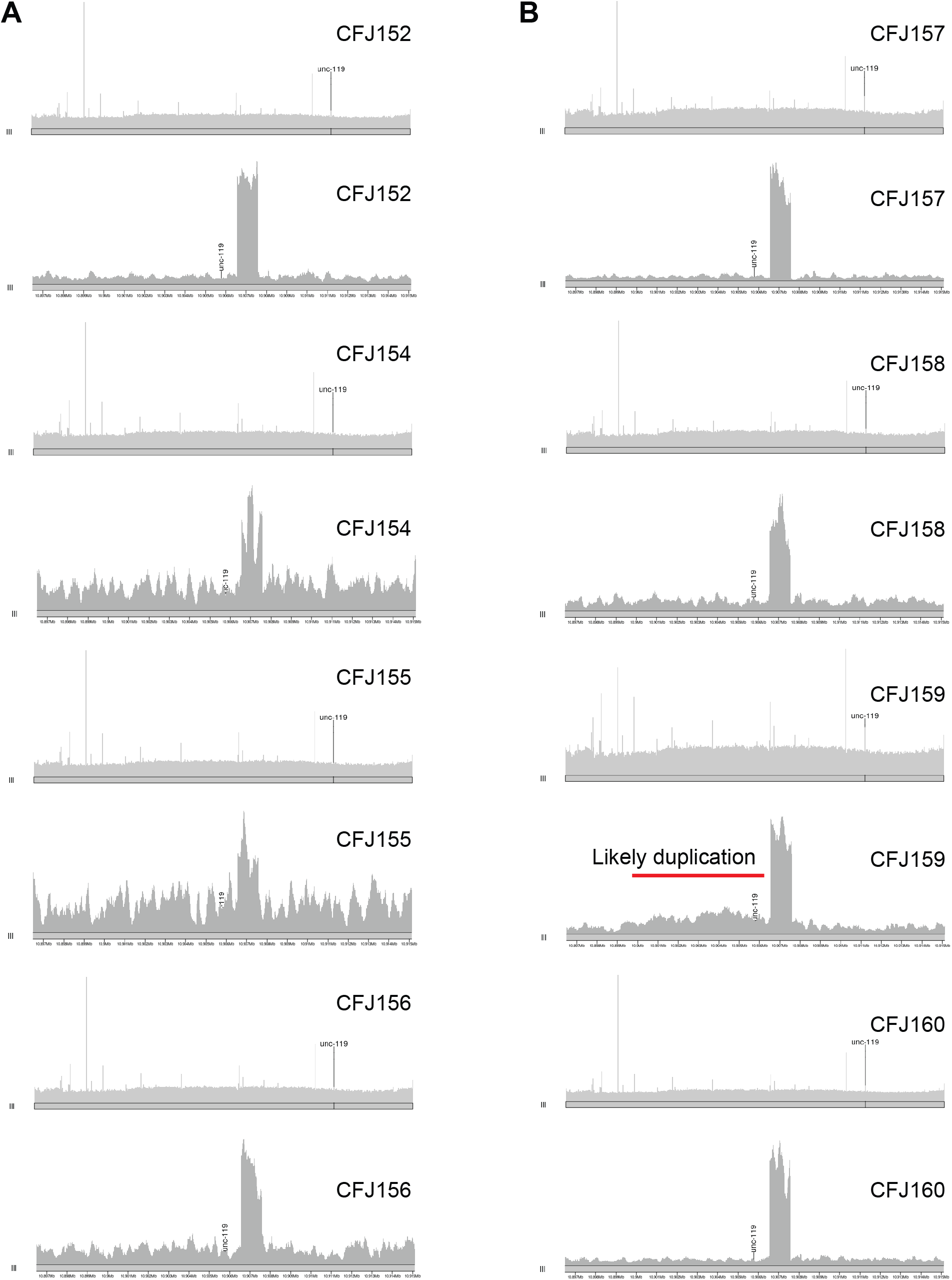
| Sequencing read coverage to detect large-scale duplications and deletions. Read coverage for eight strains carrying integrated arrays at the *unc-119*(*ed3*) locus. **A**. Array integrations generated by the “one-step plasmid” protocol. **B**. Array integrations generated by the “two-step protein” protocol. Note, the likely short duplication approximately 6-7 kb) at the *unc-119* locus in CFJ159.

**Supplementary Table S1** - Strains, plasmids, insertion sites, SNPs CFJ42 and CFJ77

**Supplementary File 1** - GenBank annotated plasmid sequence files.

